# Evidence for two distinct thalamocortical circuits in retrosplenial cortex

**DOI:** 10.1101/2021.06.11.448056

**Authors:** Eleonora Lomi, Mathias L. Mathiasen, Han Yin Cheng, Ningyu Zhang, John P. Aggleton, Anna S. Mitchell, Kate J. Jeffery

**Affiliations:** Department of Experimental Psychology, University of Oxford, The Tinsley Building, Mansfield Road, Oxford OX1 3SR, United Kingdom; School of Psychology, Cardiff University, Cardiff, UK; Department of Psychological and Brain Sciences, Dartmouth College, Hanover, New Hampshire, USA; Institute of Neuroscience, State Key Laboratory of Neuroscience, CAS Center for Excellence in Brain Science and Intelligence Technology, Chinese Academy of Sciences, Shanghai 200031, China; Institute of Behavioural Neuroscience, Division of Psychology and Language Sciences, University College London, London WC1E 6BT, UK

**Keywords:** Retrosplenial cortex, anterior thalamus, thalamocortical, theta-modulation, spatial memory, chronic rodent electrophysiology.

## Abstract

Retrosplenial cortex (RSC) lies at the interface between perceptual and memory networks in the brain and mediates between these, although it is not yet known how. It has two distinct subregions, granular (gRSC) and dysgranular (dRSC). The present study investigated how these subregions differ with respect to their electrophysiology and connections, as a step towards understanding their functions. gRSC is more closely connected to the hippocampal system, in which theta-band local field potential oscillations are prominent. We therefore compared theta-rhythmic single-unit activity between the two RSC subregions and found, mostly in gRSC, a subpopulation of non-directional cells with spiking activity strongly entrained by theta oscillations, suggesting a stronger coupling of gRSC to the hippocampal system. We then used retrograde tracers to examine whether differences in neural coding between RSC subregions might reflect differential inputs from the anterior thalamus, which is a prominent source of RSC afferents. We found that gRSC and dRSC differ in their afferents from two AV subfields: dorsomedial (AVDM) and ventrolateral (AVVL). AVVL targets both gRSC and dRSC, while AVDM provides a selective projection to gRSC. These combined results suggest the existence of two distinct but interacting RSC subcircuits: one connecting AVDM to gRSC that may comprise part of the cognitive hippocampal system, and the other connecting AVVL to both RSC regions that may link hippocampal and perceptual regions. We suggest that these subcircuits are distinct to allow for differential weighting during integration of converging sensory and cognitive computations: an integration that may take place in thalamus, RSC or both.

**Highlights:**

- The two retrosplenial cortex subregions, gRSC and dRSC, differ in their temporal firing characteristics and relation to theta oscillations.
- There are differential afferents from the anteroventral thalamic nucleus to gRSC and dRSC, with the dorsomedial subnucleus projecting selectively to gRSC.
- The anteroventral thalamus-retrosplenial cortex circuitry thus comprises two functionally and anatomically distinct but connected circuits, differentially connected to the hippocampal system, that may support the integration of cognitive and perceptual information.

## 1.0 Introduction

Retrosplenial cortex is a brain region lying between primary sensory cortices and the hippocampal spatial memory system. It has two component subregions, granular (gRSC) and dysgranular (dRSC), that may serve different, albeit related, functions (Aggleton et al., this issue). In order to shed light on these functions we first explored how gRSC and dRSC differ electrophysiologically and then traced the thalamic afferents, which form some of the most prominent inputs to RSC, to try and shed light on what may account for these differences.

Retrosplenial cortex (RSC) is a fundamental component of an extended hippocampal circuit supporting learning and memory (Vann et al., 2009). The hippocampus, RSC and anterior thalamus (ATN) also comprise core brain areas linked to neurodegeneration in Alzheimer’s disease (Braak and Braak, 1991; Raji et al., 2009). Spatial memory is a prominent component of this system: selective RSC damage in humans or animals results in memory deficits and also spatial disorientation (Reed and Squire, 1997; Sutherland and Hoesing, 1993; Vann and Aggleton, 2002, 2004, 2005; Whishaw et al., 2001; Pothuizen et al., 2008, 2009, 2010; Aggleton et al., 1995; Cooper and Mizumori, 1999, 2001). Human functional imaging studies provide further evidence of the spatial functions attributed to the RSC (Maguire, 2001; Auger et al., 2012). While rodent studies have also characterized the spatial functions of the RSC (Vann et al., 2009) the details are far from complete (Mitchell et al., 2018), although directional processing is a prominent feature (Cho and Sharp, 2001; Lozano et al., 2017; Alexander et al., 2020; Chen at al., 1994).

The topography of connections of each of the major RSC subregions, gRSC and dRSC, represents a key aspect for functional considerations on the RSC involvement in navigation and memory (Vann et al., 2009). While the RSC has critical connections with three main areas, the hippocampal formation, ATN, and visuospatial neocortical areas, these connections are not equally shared between the two RSC subregions (van Groen and Wyss, 1990, 1990b, 1992; Van Groen and Wyss, 2003; Vogt and Miller, 1983; Cenquizca and Swanson, 2007; Sripanidkulchai and Wyss, 1986; Sugar et al., 2011). gRSC connectivity is strongly coupled with regions of the hippocampal formation that are involved in spatial navigation and memory processes, as it receives direct inputs from the dorsal hippocampus, primarily from subiculum (Meibach and Siegel, 1977; Yamawaki et al., 2019; Opalka et al., 2020), and secondarily from CA1, both glutamatergic (Haugland et al., 2019; van Groen and Wyss, 1990) and GABAergic (Opalka et al., 2020; Unal et al., 2015; Jinno et al., 2007). In contrast, dRSC does not receive direct CA1 inputs and receives only sparse subicular inputs (van Groen and Wyss, 1990b, 1992; Vogt and Miller, 1983; Witter et al., 1990).

The gRSC also receives extensive inputs from basal forebrain structures that are involved in the generation of hippocampal theta (6-12Hz) oscillations. These regions are the medial septum, and diagonal and horizontal band of Broca (Unal et al., 2015; Van Groen and Wyss, 2003). Across these regions, theta oscillations are clearly present in the local field potential (LFP) and are believed to reflect the relaying of information (perhaps timing) related to hippocampal function (Burgess et al., 2002; Buzsáki, 2002, 2005; Hasselmo and Eichenbaum, 2005). By contrast, instead of these limbic inputs, dRSC is characterized by stronger connections directly from primary/secondary visual cortices and laterodorsal/ lateroposterior thalamus (van Groen and Wyss, 1990, 1990b, 1992), which are considered to endow dRSC with a privileged role in visuospatial processing (Vann & Aggleton, 2005). Jacob et al. (2017) extended this idea by showing a functional difference in the electrophysiological recordings between RSC subregions in rats. While global head direction encoding was present in both gRSC and dRSC, a directional signal more sensitive to local visual landmarks was found only in dRSC (Jacob et al., 2017). Conjunctive head direction and landmark coding in dRSC has been accounted for by reference to its stronger visual inputs as compared to global directional inputs ascending from the ATN (Page and Jeffery, 2018; Fischer et al., 2020).

Within the ATN, the anterodorsal (AD) and anteroventral (AV) nuclei are reciprocally interconnected to the two RSC subregions, although both appear to have a stronger relationship with the granular compared to the dysgranular part (Witter et al., 1990; Shibata, 1993, 1994; Van Groen and Wyss, 1995). While 56% of AD neurons are head direction cells (Taube, 1995; Taube, 2007; Clark and Taube, 2012), the majority of AV neurons (75%) have firing entrained by the theta oscillations (Vertes et al., 2001; Albo et al., 2003; Tsanov et al., 2010, 2011), suggesting that two main information streams, head direction vs theta, ascend through the ATN, with a parallel organization (AD vs AV). Given that theta propagation is thought to serve spatial mnemonic functions of the limbic system by optimizing synaptic plasticity and inter-regional communication (Hyman et al., 2003; Buzsáki, 2002, 2005; Buzsáki and Moser, 2013), we used theta as a marker for hippocampal coupling to determine whether gRSC and dRSC have distinct interactions with hippocampus. We hypothesized that theta-rhythmic firing would be stronger in gRSC compared to dRSC, reflecting a stronger interaction with the hippocampal navigational network. We then used retrograde tracing to see if the anterior thalamic inputs to the two subregions could be differentiated. As we show here, we found that gRSC is more theta-modulated than dRSC, and with anatomical tracing we then found that it selectively receives a stronger input from dorsomedial AV (AVDM). These findings strengthen the evidence that gRSC is part of a subcircuit that connects with the hippocampal system, while dRSC is a parallel circuit that interacts more strongly with sensory cortices. We finish with some speculations about a possible role for RSC in linking these two systems.

## 2.0 Methods and Materials

### 2.1 Electrophysiology recording experiment

#### 2.1.1 Animals and coordinates

For this study we used data from four male Lister Hooded rats (Charles River, UK) that were surgically implanted with tetrodes. Two of the rats (R602, R636) had participated in a previously published experiment (Jacob et al., 2017) and two were part of a new experiment (Zhang et al., in prep). We only included rats in which we could identify the electrode track in the histological section, and for which histology indicated that electrodes sampled both RSC subregions (electrodes advanced from dRSC into gRSC).

Surgery followed standard stereotaxic surgery techniques for chronic recording of extra-cellular activity in behaving animals (Szymusiak and Nitz, 2003). Implant coordinates were based on previous study of head direction cells in the RSC of Jacob et al. (2017). These were as follows: AP, -5.5; -5.8; ML, ±1.0 - ±0.80; DV, 0.4 - 0.65 (in mm from Bregma). All rats were implanted in the left hemisphere. All experiments were carried out in accordance with UK Animals (Scientific Procedures) Act, 1986 and EU directive (2010/63/EU), complying with ARRIVE guidelines for the care and use of laboratory animals.

#### 2.1.2 Behavioral protocol and testing apparatus

Animals were screened daily for single unit activity. Screenings were made in an open field box in within a cue-rich room. When a set of units was isolated, the animal was transferred to the experimental room and the recording session was conducted in a two-compartment box, which was configured to test a hypothesis (not addressed here) about directional encoding. Briefly, the apparatus comprised two identical rectangular compartments connected by a central doorway in the long wall that separated them (for details see Jacob et al., 2017). One short wall of each compartment was adorned with a large white cue card, and the compartments were differentially scented, one with lemon and one with vanilla. The apparatus was located within a curtained-off area. Each recording session consisted of five consecutive trials, with each trial involving a different box orientation. To promote exploratory behavior throughout the recording, rice was randomly thrown in the box by the experimenter. For the purposes of the present analysis, only the first 10min-long baseline trial was considered.

#### 2.1.3 Signal processing and tracking

Single unit and LFP signals were recorded using a headstage amplifier connected to a microdrive and connected to the multichannel recording system (DacqUSB, Axona) via a flexible lightweight cable. Single unit activity was continuously sampled (48 kHz sampling frequency), amplified 15-50K times and band-pass filtered from 300 Hz to 7 kHz. The signal from each electrode was differentially recorded against the signal from an electrode on a different tetrode with low spiking activity. Spikes were defined as short-lasting events exceeding a user-defined voltage threshold. Spikes were time- stamped and stored together with a 1ms of surrounding signal (with 200 µs pre- and 800 µs post-threshold interval). Tetrodes were lowered by 25-150 µm to locate single-unit spiking activity. The LFP signal was collected single-ended (250 Hz sampling frequency), amplified 5k times, filtered with a 500 Hz low-pass filter and stored with the unit data. A 50 Hz notch filter was applied to remove local mains noise. Two light-emitting diodes (LEDs), one large and one small and separated by 5 cm, were attached to the headstage to provide the position and the orientation of the rat’s head. The LEDs were imaged with an overhead camera (50 Hz sampling rate). Spike times, LFP signal, position in x- and y-coordinates and directional heading in degrees were saved for offline analysis.

#### 2.1.4 Data analysis

Data were analyzed using TINT (Axona, UK) and MATLAB R2018a (MathWorks, USA) with custom-made programs and functions. CircStat toolbox was used for analysis using circular statistics (Berens, 2009).

#### 2.1.5 Spike sorting

Spike-sorting was performed using KlustaKwik (Kadir et al., 2014) followed by manual refinement using the TINT software package (Axona). To confirm that isolated clusters resulted from single unit activity, the following inclusion criteria were adopted: (1) an average waveform displaying the classical action potential shape, and (2) fewer than 10% of spikes occurring in the first 2ms of the autocorrelogram (action potential refractory period). Only cells meeting these criteria were accepted into further analysis.

#### 2.1.6 Spectral analysis

The LFP signal, recorded by low-pass filtering the raw neural signal, was analyzed using a fast Fourier transform (FFT). The power of each frequency component was defined as the square of the magnitude of the output of the FFT. To compare across animals, power measures within each trial were normalized to z-scores and then plotted and smoothed with a Gaussian kernel (bandwidth = 2Hz; standard deviation = 0.5Hz). The power of theta frequency was found as the maximum z-score within the movement-associated (Type-1) theta frequency range (6-12Hz). Average theta frequency was found as the frequency value associated to peak theta power.

#### 2.1.7 Theta analysis

To investigate spike-LFP coupling, the LFP signal was filtered offline for Type-1 theta using a 4^th^ order Butterworth filter with band-pass between 6-12Hz. A zero-phase filtering was applied to prevent the phase shift introduced by the Butterworth filter. The Hilbert transform was then applied to the bandpass-filtered LFP to derive instantaneous theta phases (250Hz sampling rate). Each theta cycle was defined such that peaks occurred at 0^◦^ and 360^◦^, with a trough at 180^◦^. To determine at which point to apply the Hilbert Transform, the LFP signal (Hz) was linearly interpolated with spike time (s), with each spike being assigned to the phase of the theta cycle at which it occurred. To visualize the spike-LFP relationship, we then double-plotted a spike-phase histogram for each cell. The probability distribution of spikes relative to theta phases was derived by smoothing the spike-phase histogram using a circular kernel density estimation (KDE) method, with an automatically selected bandwidth parameter using the plug-in rule of Taylor (2008).

Theta spike-LFP coherence and rhythmicity were assessed by computing an index of theta phase-coupling (IC) and an index of rhythmicity (IR), respectively. Coupling refers to the phase of theta at which spikes occurred, as spikes may be emitted with a timing coupled to theta (phase-locked) even if there is no overt rhythmicity on the autocorrelogram (Eliav et al., 2018). We computed an IC by computing the mean vector length (Rayleigh vector, CircStat toolbox) of the spikes vs theta phase distribution (Climer et al., 2015; Frank et al., 2001). For each cell, spike phases were binned between 0 and 360^°^ with 6^°^ bins. For each cell with IC > 99^th^ percentile of shuffle distribution, the preferred theta phase was estimated as the circular mean of all the spike phases, and a Rayleigh test was conducted to assess the significance of this phase preference across cells (see Supplementary materials, Section 1.2). This value was used to sort cells into theta-modulated vs. not.

The IR additionally quantifies the temporal modulation of a cell’s firing at theta frequency range (6-12Hz). A theta-rhythmic cell will show peaks at ∼125ms on its autocorrelogram. Autocorrelograms of the spike trains were plotted between ± 500ms using 10ms bins, normalized to the maximum value and smoothed (20 bins boxcar). Following methods outlined in Lozano et al. (2017), the IR was calculated as the difference between the expected theta modulation-trough (autocorrelogram value between 60-70ms) and the theta-modulation peak (autocorrelogram value between 120-130ms), divided by their sum. It takes values between -1 and 1.

#### 2.1.8 Cell identification and inclusion criteria

Head direction and bidirectional cells were identified as described in Jacob et al. (2017), and removed from the current analysis, as they were previously included in Jacob et al. (2017), and they were not theta-modulated. Briefly, to analyze the directional firing characteristics of single units, the spike times and position samples of the rat’s facing direction were sorted into bins of 6°. The mean firing rates per angular bin (Hz) were smoothed with a 5-bin (30°) smoothing kernel and polar-plotted. Finally, the angles were doubled and plotted modulo 360° in order to remove any bidirectionality that might be present in the tuning curves of bidirectionally tuned cells.

To be considered for analysis, cells needed to have (1) a tuning curve that was not obviously unimodal/bidirectional from visual inspection; (2) Rayleigh vector length of the doubled head direction angles < 0.22, which is the original threshold used by Jacob et al. (2017) and (3) average firing rate > 0.1Hz for the trial.

To provide a control for each cell against which to measure its theta-phase coupling we created a time-shifted null distribution in which the entire sequence of spike times was circularly time-shifted by a random amount lying between 20s and the duration of the recording session minus 20s. This preserved the number of spikes and the temporal structure of the spiking sequence, but dissociated the time of spiking from the theta oscillations. This was repeated 1000 times for each cell, and the IC was computed for each repetition. All values were pooled together to derive a control distribution, representing the chance level for which the activity of each RSC cell was locked to theta. A cell was considered as significantly modulated at theta frequency if the IC was >99^th^ percentile of the control distribution for that neuron.

#### 2.1.9 Cell-specific temporal firing characteristics

To investigate bursting properties of RSC neurons, inter-spike intervals (ISI) were plotted in a frequency histogram between 0-250ms. Cells with peak ISI <10ms, which defines a burst mode (Fanselow et al., 2001), were classified as bursting and the remainder as tonic. The burst index was calculated as the ratio of spikes with peak ISI <10ms to all the spikes in a trial (Mizuseki et al., 2012). Burst index values closer to 1 indicate the presence of intra-burst spikes, whereas smaller values indicate a prevalence of single spikes.

#### 2.1.10 Waveform analysis

Peak-to-trough waveform width was used to separate narrow from broad spiking cell types (Lewicki, 1998). We followed methods reported in Ardid et al. (2015), using the open source code from the public Git repository: https://bitbucket.org/sardid/waveformanalysis (Waveform Analysis toolbox; Ardid et al., 2015). Briefly, a two-Gaussian model was applied to the distribution of waveform widths. We defined two cutoffs on this model that divided waveforms into three groups: narrow, broad and in-between. Broad waveforms were those for which the likelihood to be considered broad was larger than 10 times the likelihood to be narrow. Narrow waveforms were those for which the likelihood to be considered narrow was larger than 10 times the likelihood to be broad. Unclassified waveforms were those falling into the dip of the bimodal distribution. The bimodality of the distribution was tested using the calibrated version of the Hartigan dip test (*p* < 0.01).

#### 2.1.11 Statistics

For gRSC-dRSC comparisons, if data were found to deviate significantly from a normal distribution (Matlab functions *kstest*), we used a non-parametric test (Wilcoxon Rank-sum test, WRS, Matlab function *ranksum*). Otherwise, we used parametric tests. To evaluate simultaneously the effect of two grouping variables (brain region and cell type) on waveform parameters, we used a two-way ANOVA. We also used the Chi-squared goodness-of-fit test (*X*^2^) when testing observed numbers of theta-rhythmic cells in gRSC and dRSC against an expected equal proportion, and the number of interneurons against an expected equal proportion. To test the significance between two circular distributions, we employed the Watson-Williams multi-sample F-test for equal circular means (WW, Matlab function *circ_wwtest*, Circular statistics toolbox), in addition to the Rayleigh test. All statistical tests are two-tailed. For all box plots, a line is marked at the sample median, boxes at the interquartile range, whiskers at the minimum and maximum value. Markers represent individual data points. In all figures and tables, *p*-value significance level: *∗* = 0.05, *∗∗* = 0.01, *∗ ∗ ∗* = 0.001.

### 2.2 Retrograde tracer experiment

#### 2.2.1 Animals and tracers

Twenty-three male Lister hooded rats (Charles River, UK; Envigo, UK) weighing 291-614g at the time of surgery were used for the retrograde tracer experiments. Four retrograde tracers were employed for a total of 29 injections: (i) cholera toxin subunit B (CTB; Polysciences Inc, Eppelheim, Germany), (ii) fast blue (FB; Polysciences Inc, Eppelheim, Germany), (iii) biotinylated dextran amine tracer (BDA; 3 kD version; Life Technologies Ltd, Paisley, UK), and (iv) fluorogold (FG; Fluorochrome Inc., Denver, CO, USA). The CTB was used either unconjugated or conjugated with Alexa Fluor 594 (Ctb-AF594; Molecular Probes, Invitrogen Inc.) or with Alexa Fluor 488 (Ctb-AF488; Molecular Probes, Invitrogen Inc.). The single BDA case was also included in a previous study (Mathiasen et al., 2017), which provides full methodological procedures for this tracer. Six animals had dual injections while the others had single injections placed in either gRSC or dRSC.

#### 2.2.2 Surgery

All injection surgeries took place under isoflurane anesthesia (isoflurane-oxygen mixture 1.5-2.5%) with the rat positioned in a stereotaxic frame (Kopf Instruments, Tujunga, CA). A dorsal craniotomy exposed the central sinus and adjacent tissue. Lambda and bregma were set at the same depth coordinates, creating a flat skull. Tracers were injected either mechanically (all FB, FG, Ctb-AF594, Ctb-AF488 injections) via a 0.5µl, 0.66µl or 1.0µl Hamilton pipette (Hamilton, Bonaduz, Switzerland) or iontophoretically. See Supplementary materials **(Tables 3-4)** for injection coordinates. Mechanical tracer injections were infused with a flow of 10-20 nl/min. In all cases, the needle was left in place for a further 10min before being retracted. As a part of the analgesia regime, Lidocaine was administered topically to the scalp (0.1ml of 20 mg/ml solution; B. Braun, Melsungen, Germany) and meloxicam was given subcutaneously (0.06ml of 5mg/ml solution, Boehringer Ingelheim Ltd, Berkshire, UK). After 6-8 days post-injection, each rat was perfused and histological analysis was performed (see section Histology).

### 2.3 Histology and tracing

At the completion of all experiments, rats received an overdose of sodium pentobarbital (Euthatal, Merial, Harlow, United Kingdom) and were perfused transcardially with 0.1 M phosphate-buffered saline (PBS) followed by paraformaldehyde (PFA) perfusion (4% paraformaldehyde solution in 0.1 M PBS) or with saline (0.9% sodium chloride solution) followed by 10% formalin perfusion (10% formalin in 0.9% sodium chloride solution). The brains were removed, stored in 10% formalin or PFA, followed by 25-30% sucrose solution (25-30% in 0.1 M PBS), and finally frozen with dry ice. Sections were cut coronally (30-50µm) with a freezing microtome or cryostat and were mounted on glass slides. Tetrode locations were histologically verified from cresyl violet-stained brain section. Digital pictures were acquired with an Olympus microscope (Olympus Keymed, Southend-on-Sea, U.K.)

In cases with unconjugated CTB tracer injections, sections were first washed for 3x10 min in 0.1 M PBS, washed 3 x 10 min in PBS-TX (0.2% Triton X-100 in 0.1M PBS) and incubated with the primary antibody rabbit anti-CTB overnight (1:3000) (Sigma-Aldrich, UK). Sections were washed for 3 x 10 min (PBS-TX) and incubated with the secondary antibody goat anti-rabbit (1:200) (conjugated to DyLight 594; Vector Laboratories, CA) for two hours. After a further 3 x 10 min wash (PBS), sections were mounted. Sections were then further dehydrated (ethanol), defatted (xylene), and coverslipped with DPX. All slides were mounted in DPX or Vectashield antifade mounting medium (Vectorlab, H-1000). Slides from animal 717 were mounted in Vectashield antifade mounting medium with propidium iodide (Vectorlab, H-1300) which stained the nucleus red, while slides from animal 878 were mounted in Vectashield antifade mounting medium with DAPI (Vectorlab, H-1200) which stained the nucleus blue. Slides were analysed for retrograde cell labelling in the AV, using epi-fluorescence microscopy for CTB, FG and FB labelling and brightfield microscopy for the single case with BDA label. For visualization purposes, FG labelled images were pseudo-coloured in cyan, CTB and Ctb-AF594 in red, Ctb -AF488 in green and FB in blue.

## 3.0 Results

### 3.1 RSC electrophysiological experiment

#### 3.1.1 Histology

As mentioned earlier, for the analysis we only included four rats for which histology indicated that electrodes sampled both RSC subregions (electrodes advanced from dRSC into gRSC). For each electrode bundle, we marked on the relevant histological section our best estimate of the electrode tip. The cell locations were estimated based on the reconstruction of the electrode tract from histology and the record of screw turns that lowered the electrodes each day, starting from a known DV within the dRSC. **Figure 1** illustrates examples of brain sections in which the electrode track is clearly visible. For one animal (R887), the tract suggested that electrodes were implanted more laterally and were advanced through the white matter tract of the cingulum fibres (Cg; **Figure 1;** See Supplementary materials, **Section 1.1)** and this could explain the under sampling of cells from this animal (see **Table 1,** first row).

**Figure 1.**
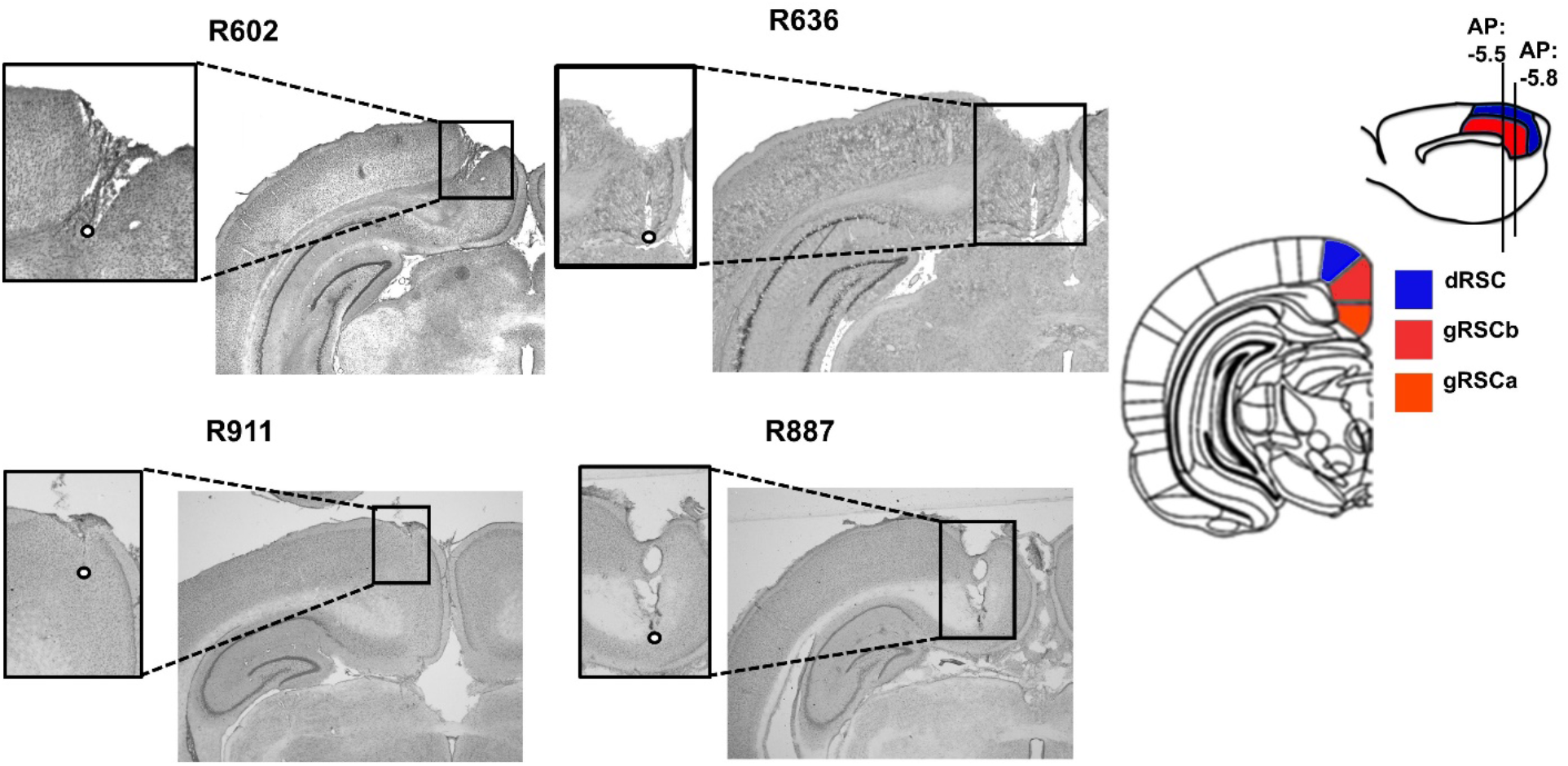
Histology of RSC recordings. Left: Photomicrographs of the Nissl-stained coronal sections through the recording site, for each of the four rats implanted in RSC (left hemisphere). White dots depict the final electrode depth. Top right: schematic of the AP-level (mm) of the recording sites within the RSC. R636 and R602, AP = -5.8; R911 and R887, AP = -5.5. AP-levels estimated based on the shape of cingulum fibers (Cg) and the width of the central sulcus (See supplementary materials, Section 1.1.). Middle right: schematic of the RSC subregions for a coronal AP level of -5.5, adapted from Paxinos and Watson (2007). Images for R602 and R636 were supplied by the authors of Jacob et al. (2017).

**Table 1.**
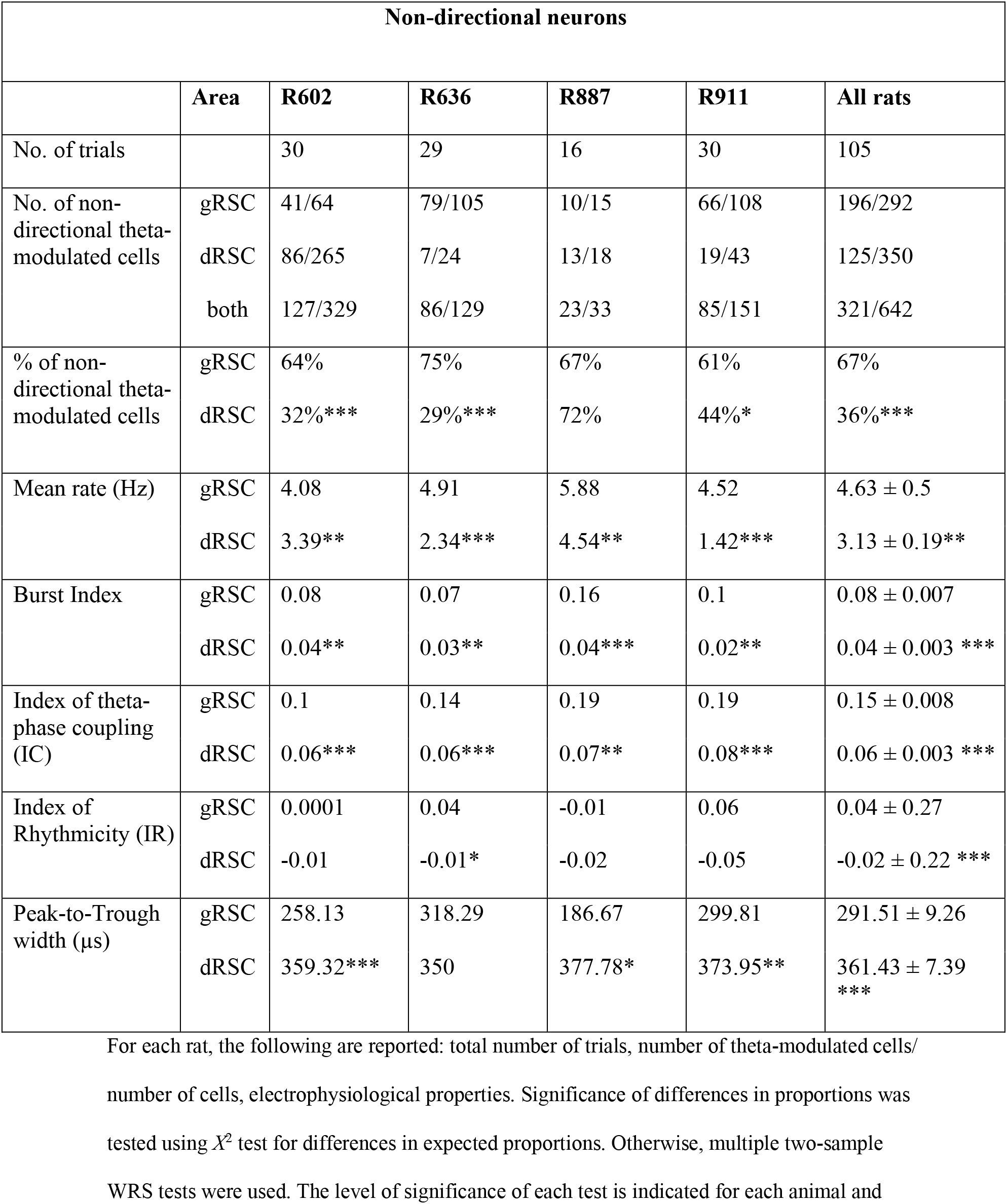
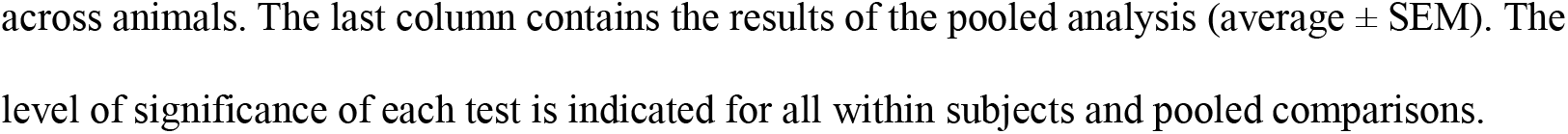
Distribution and electrophysiological properties of non-directional RSC neurons. Average for each animal and combined across animals, compared between gRSC and dRSC.

| Non-directional neurons |  |  |  |  |  |  |
| --- | --- | --- | --- | --- | --- | --- |
|  | Area | R602 | R636 | R887 | R911 | All rats |
| No. of trials |  | 30 | 29 | 16 | 30 | 105 |
| No. of non-directional theta-modulated cells | gRSC | 41/64 | 79/105 | 10/15 | 66/108 | 196/292 |
|  | dRSC | 86/265 | 7/24 | 13/18 | 19/43 | 125/350 |
|  | both | 127/329 | 86/129 | 23/33 | 85/151 | 321/642 |
| % of non-directional theta-modulated cells | gRSC | 64% | 75% | 67% | 61% | 67% |
|  | dRSC | 32%*** | 29%*** | 72% | 44%* | 36%*** |
| Mean rate (Hz) | gRSC | 4.08 | 4.91 | 5.88 | 4.52 | 4.63 ± 0.5 |
|  | dRSC | 3.39** | 2.34*** | 4.54** | 1.42*** | 3.13 ± 0.19** |
| Burst Index | gRSC | 0.08 | 0.07 | 0.16 | 0.1 | 0.08 ± 0.007 |
|  | dRSC | 0.04** | 0.03** | 0.04*** | 0.02** | 0.04 ± 0.003 *** |
| Index of theta-phase coupling (IC) | gRSC | 0.1 | 0.14 | 0.19 | 0.19 | 0.15 ± 0.008 |
|  | dRSC | 0.06*** | 0.06*** | 0.07** | 0.08*** | 0.06 ± 0.003 *** |
| Index of Rhythmicity (IR) | gRSC | 0.0001 | 0.04 | -0.01 | 0.06 | 0.04 ± 0.27 |
|  | dRSC | -0.01 | -0.01* | -0.02 | -0.05 | -0.02 ± 0.22 *** |
| Peak-to-Trough width (μs) | gRSC | 258.13 | 318.29 | 186.67 | 299.81 | 291.51 ± 9.26 |
|  | dRSC | 359.32*** | 350 | 377.78* | 373.95** | 361.43 ± 7.39 *** |

For each rat, the following are reported: total number of trials, number of theta-modulated cells/ number of cells, electrophysiological properties. Significance of differences in proportions was tested using $\chi^2$ test for differences in expected proportions. Otherwise, multiple two-sample WRS tests were used. The level of significance of each test is indicated for each animal and
across animals. The last column contains the results of the pooled analysis (average ± SEM). The level of significance of each test is indicated for all within subjects and pooled comparisons.

#### 3.1.2 The same LFP theta oscillations are recorded from gRSC and dRSC

Consistent with previous work in freely moving animals, the LFP showed characteristic network patterns, including clear theta oscillations (Dillingham et al., 2019; Young and McNaughton, 2009; Borst et al., 1987; Talk et al., 2004). Theta oscillations from gRSC and dRSC did not differ in average peak Type-1 (movement-associated; Olvera-Cortés et al., 2002; Caplan et al., 2003) theta power and frequency (6-12Hz; **Figure 2**). Average peak theta power was 6.88 *±* 0.38 for gRSC recordings and 6.577 *±* 0.35 for dRSC recordings (WRS test, z = 0.7, *p* = 0.49). The associated average frequency was 8.73 *±* 0.14Hz for gRSC recordings and 8.63 *±* 0.16Hz for dRSC recordings (z = -0.84, *p* = 0.40). The lack of differences between gRSC and dRSC LFP was maintained when the data were segregated according to animal (WRS test, all *p* > 0.05).

**Figure 2.**
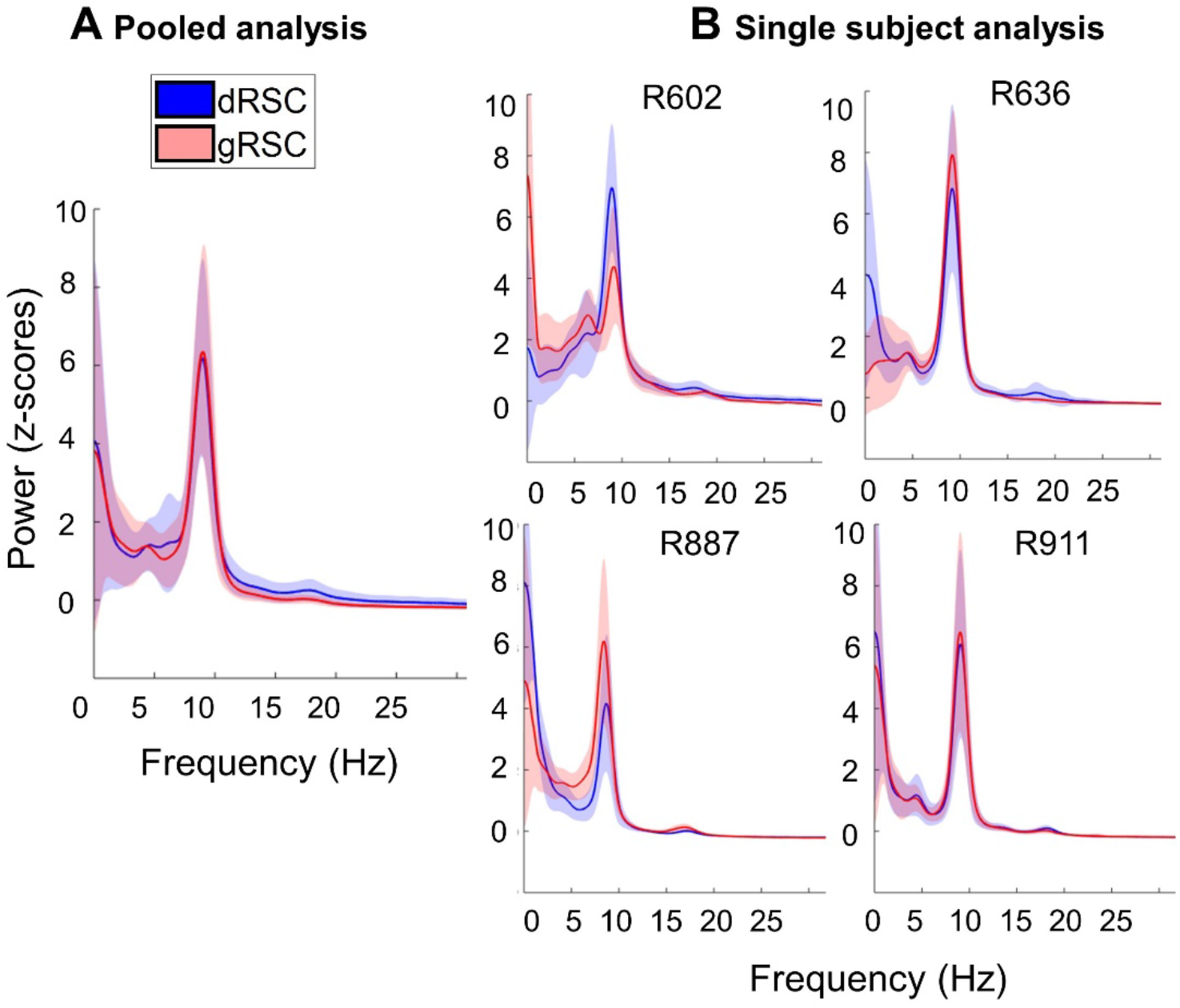
Theta oscillations have the same power and frequency. Average normalized power spectrum across all trials, **(A)** combined across all animals (n=4; pooled analysis), and **(B)** within each animal (single subject analysis). Shaded area is the standard deviation. X-axis, frequency (Hz); y-axis, power (z-scores). No difference in LFP theta oscillations were seen between gRSC and dRSC.

#### 3.1.3 An anatomically organized theta-modulated cell population exists in the RSC

We then looked at theta-band properties of spiking in single neurons. A total of 642 non-directional cells were analyzed from the four rats moving freely in the two-compartment box to determine whether there is differential coupling of gRSC and dRSC with the hippocampal navigational network, reflected in differential theta modulation. Single-cell results were obtained first after pooling cells across animals into a single distribution (pooled analysis), and second, from cells recorded for each animal individually (single subject analysis). Unless otherwise stated, the same results were observed in both analyses and between animals.

We first assessed whether cells were theta-modulated by determining their IC. Of the 642 non-directional cells identified, 292 cells came from gRSC and 350 from dRSC. Of these cells, half (n = 321/642, 50%) reached the criterion for significant theta modulation, based on the IC being greater than 99^th^ percentile of the control distribution for that neuron. **Table 1** reports a summary of recording locations and number of theta-phase-locked cells recorded in each RSC subregion for the four animals.

Despite the lack of LFP theta differences between gRSC and dRSC recordings discussed above, the distribution of theta-modulated cells varied considerably between RSC subregions, as indicated by the significance of a *X*^2^ test for differences in proportions. Across all animals, 67.1% (196/292) of gRSC cells fired in phase with theta, compared to only 35.7% (125/350) of dRSC cells (*χ*^2^ = 62.82; *p* < 0.0001). Significant differences were confirmed when the data were segregated according to animals (*X*^2^ test, all *p* < 0.05), except for one animal (R887) for which differences did not reach significance (*χ*^2^ = 0.12; *p=* 0.73; **Table 1**). As already noted, for R887 there was under-sampling of gRSC cells (n=15 across 7 trials), possibly due to the tract trajectory passing through white matter tracts before reaching gRSC. We concluded that while the majority of gRSC non-directional cells were theta-modulated, the majority of non-directional cells in dRSC fired uncoupled from theta (see pie chart in **Figure 6)**.

Theta-strength (spike-LFP coupling) and rhythmicity (autocorrelograms) were assessed by computing IC and IR, respectively (**Table 1**). At the single cell level, the average IC was significantly higher in gRSC compared to dRSC (IC: gRSC (292), 0.15 *±* 0.01; dRSC (350), 0.06 *±* 0.003; WRS test, z = -10.09, *p* < 0.001). This reflects the overall larger proportion of theta-modulated neurons in gRSC (**Figure 6**), as well as the stronger theta modulation-depth within each of these neurons. In this respect, the firing of cells that reached the theta-modulation criterion in gRSC was considerably more phase-locked to theta compared to the same cells recorded in dRSC, which only displayed weak spike-LFP coherence (IC: gRSC (196), 0.2 *±* 0.01; dRSC (125), 0.1 *±* 0.01; WRS test, z = 7.17, *p* < 0.0001; **Figure 3**). Significant IC differences were confirmed when the analysis was performed individually for each animal and no difference was observed between animals (**Table 1;** See **Supplementary Figure 1** for phase preference analysis of the theta-rhythmic neurons).

**Figure 3.**
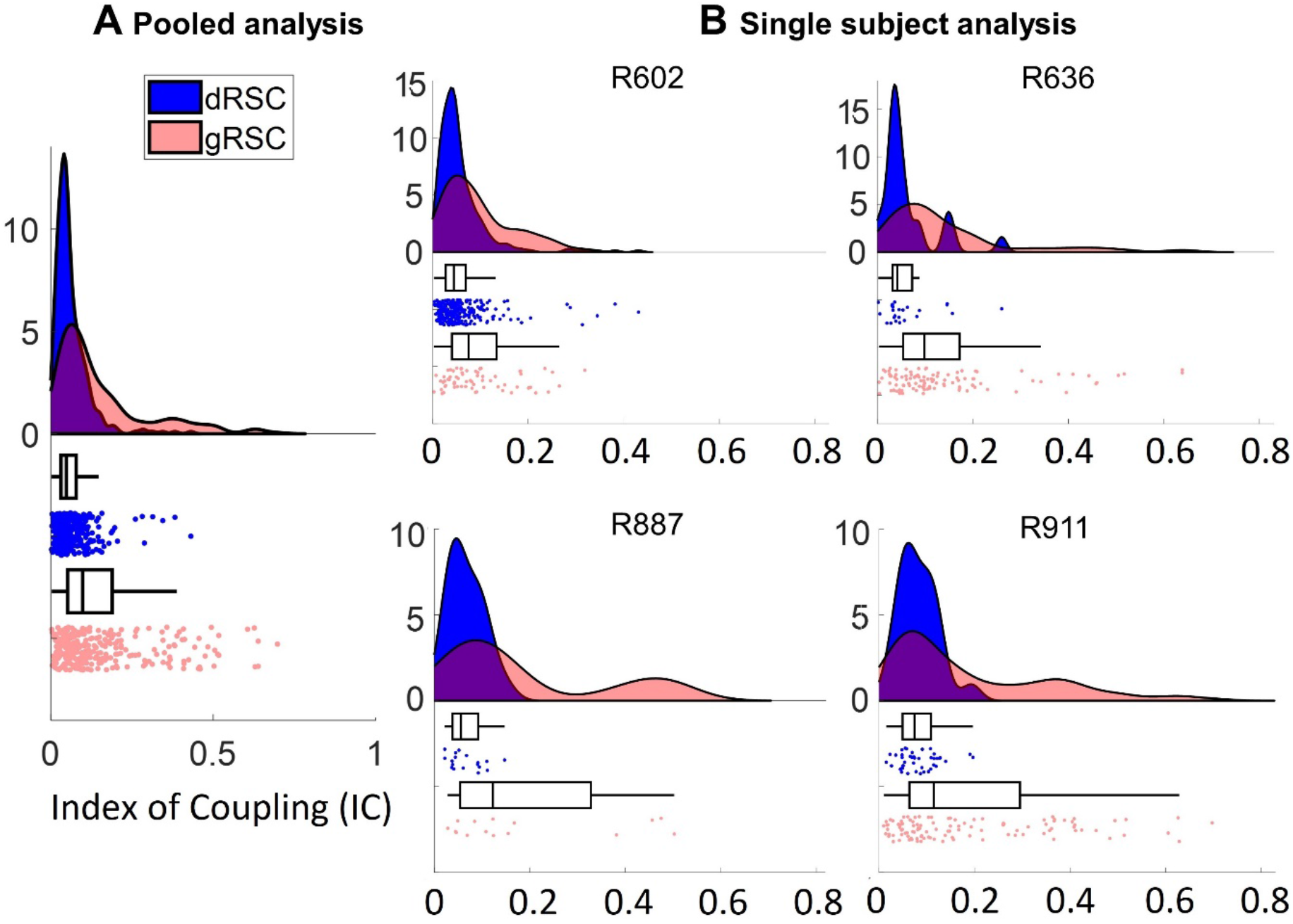
Differences in theta-phase locking. Distribution of ICs **(A)** for all cells and **(B)** within each animal. (C) Distribution of IC considering only cells with firing significantly coherent with theta (IC>99^th^ percentile shuffling). Data is shown as a split violin plot + raw jittered data point + boxplot. Split violin plots show the probability density function (PDF) of the observed data points. Each marker is a cell (dRSC, blue; gRSC, red). gRSC: the higher average IC reflects a larger proportion of neurons with firing locked to theta, as well as stronger theta-modulation depth within each neuron.

Average IR was significantly higher and more positive in gRSC compared to dRSC when all cells were pooled into a single distribution (IR: gRSC, 0.04 *±* 0.02; dRSC, -0.02 *±* 0.01; WRS test, z = -4.61, *p* < 0.0001). In the single subject analysis, however, differences in IR did not reach a statistical significance in more than half of the experimental animals (3/4; only for R636; z = -2.09; *p* = 0.04; **Table 1**), albeit in all cases averages were higher in gRSC compared to dRSC. This is because some neurons (total, 49.53%; gRSC, 42.86%; dRSC, 60%) with significant phase-locking to theta did not show significant theta rhythmicity in their spike trains, suggesting spikes emitted at theta frequency for only brief periods of otherwise theta-unrelated spikes. This discrepancy, in which more cells fired phase-locked to theta than fired rhythmically, means that our classification criteria (based on IC) is sensitive to low amounts of locking, which can potentially occur in cells with non-rhythmical spiking, as previously reported for RSC (Eliav et al., 2018; Alexander et al., 2018) and ATN neurons (Albo et al., 2003). **Figure 4** displays examples of a theta phase-locked and rhythmic cell (high IC, IR>0.001), and a theta-locked but non-rhythmic cell (high IC, IR<=0.001). As can be seen, the main difference between the two cell types, aside from the autocorrelation differences (sinusoidal vs flat), is the degree of theta modulation and height of the baseline firing (firing at low-excitability phases) that is visible in the spike-phase histograms (iii). Theta phase-locked and rhythmic cells exhibit strong locking to the theta cycle and low baseline firing outside of a narrow oscillatory range, whereas phase locked and non-rhythmic cells exhibit weaker, and sometimes virtually no locking to theta and a higher baseline firing.

**Figure 4.**
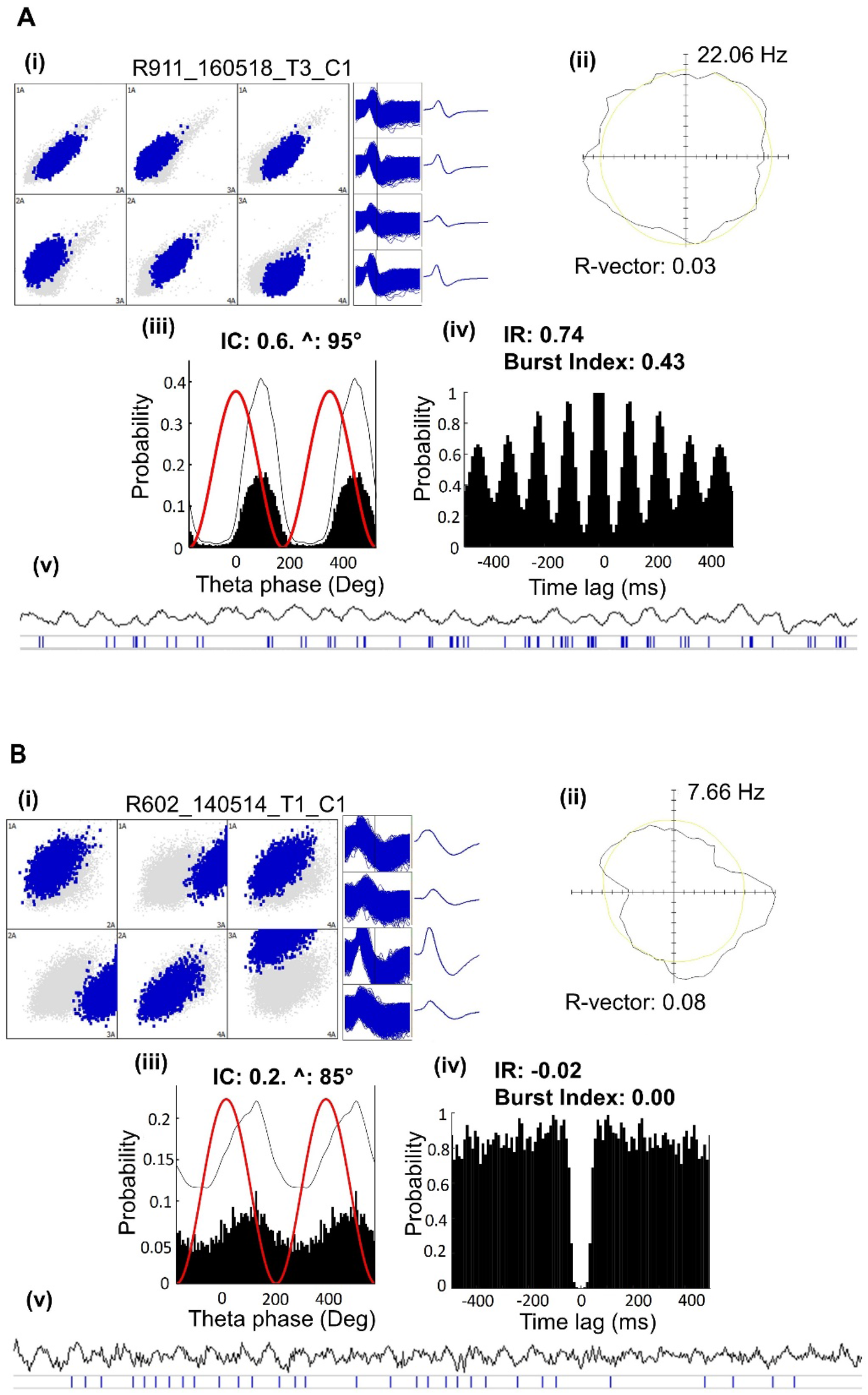
Two types of theta-modulated cells: (A) theta coupled and rhythmic vs (B) theta coupled and non-rhythmic. For each cell the following plots are shown, **(i)** Cluster space plots and associated tetrode spike waveforms. **(ii)** Polar plot of firing rate vs head direction, showing a uniform directional firing distribution (Hz). Peak firing rate is reported next to each plot. Thinner line circle indicates dwell time distribution (seconds). **(iii)** Double-plotted histogram of spike frequency relative to theta phases. Thin line is the corresponding smoothed density estimates (see Methods). The thicker line is the sine wave representing two theta cycles. For cell (B), the spike-phase histogram displays weak locking and elevated baseline firing. **(iv)** Temporal autocorrelogram showing theta-frequency rhythmicity for (A) but non-rhythmical firing for (B). **(v)** Spike raster and corresponding unfiltered LFP signal recorded from the same tetrode. Each trace shows 3s of recordings. Cell (A) fires rhythmic spike trains, roughly corresponding to the frequency of LFP. In contrast, theta-modulation is not visually detectable in the LFP traces of (B), and here is based on IC > 99^th^ percentile shuffling.

#### 3.1.4 gRSC cells have higher firing rate and spike bursting propensity

In addition to differences in the intrinsic oscillation of single cells, gRSC and dRSC displayed a number of other interesting differences in their temporal firing properties. gRSC cells fired with significantly higher mean firing rates compared to dRSC (mean firing rate: gRSC, 4.63 *±* 8.61Hz; dRSC, 3.13 *±* 3.6Hz; WRS test, z = 2.73, *p* < 0.01). Furthermore, gRSC cells displayed a higher propensity to burst. In gRSC, 88/292 cells (30%) fired in bursts with peak ISI <10ms, compared to only 41/350 cells (11.7%) in dRSC (*χ*^2^ = 33.65; *p* < 0.0001). The prevalence of bursting units in gRSC was reflected in a significantly higher average burst index (burst index: gRSC, 0.08 *±* 0.01; dRSC, 0.04 *±* 0.01; WRS test, z = -6.30, *p* < 0.001). **Figure 5** shows examples of bursting and non-bursting units. Among bursting units, bursts were discharged rhythmically within theta frequencies in gRSC, but not dRSC where cells mostly burst at other frequencies (theta-bursting cells, gRSC, 76/88, 86.3%; dRSC, 17/41, 41.1%; *χ*^2^ = 28.03; *p* < 0.0001). Hence, theta-modulated cells were further subdivided into theta-modulated non-bursting cells (gRSC, n = 120 vs dRSC, n = 108), and theta-bursting cells (gRSC, n = 76 vs dRSC, n = 17). Theta-modulated cells emitted compact spike trains at theta frequency, without bursts, as their ISI reached values >10ms. In contrast, theta-bursting cells fired rhythmically at theta frequency with ISI <10ms. Differences in firing frequency and burst index were maintained when data were segregated according to animal, and again no difference was found between animals (all *p* < 0.05; **Table 1).** Overall, these results confirm gRSC as a locus of theta-bursting neurons (**Figure 6**).

**Figure 5.**
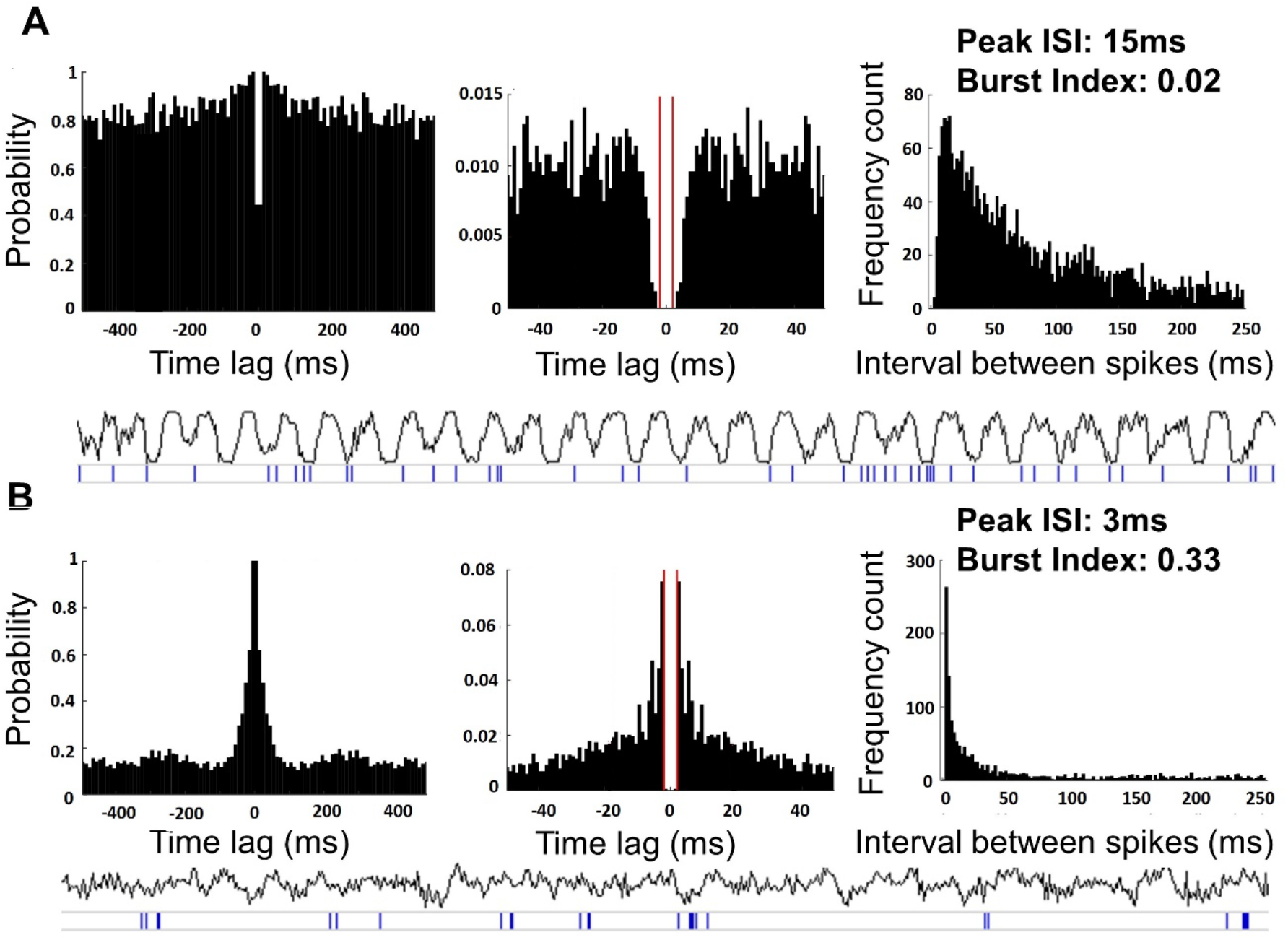
Two types of temporal firing modes: **(A)** non-bursting vs **(B)** bursting. Two examples of non-theta-modulated cells. The following plots are shown. From left to right: (1) Normalized temporal autocorrelogram implemented between ±500ms. (2) Temporal autocorrelogram implemented between ±50ms: gaps at intervals < 2ms reflect the refractory period; (3) Inter-spike interval (ISI) histogram and (4) plots of spiking events (bottom) relative to the raw LFP trace (top). Non-bursting unit (A): fires single spikes (in groups of 1-2) and displays a prevalence of long ISI. Bursting unit (B): spikes are emitted in groups of 2-3 spikes separated by ISI < 10ms, defined as a compact spike train. The prevalence of short ISI is reflected in a fast-decaying central peak of the ISI histogram and autocorrelogram, which can be contrasted to the less steep histogram peak in (A). Each spike train typically has a pause longer than 120ms before and after it, as firing is not aligned to theta.

**Figure 6.**
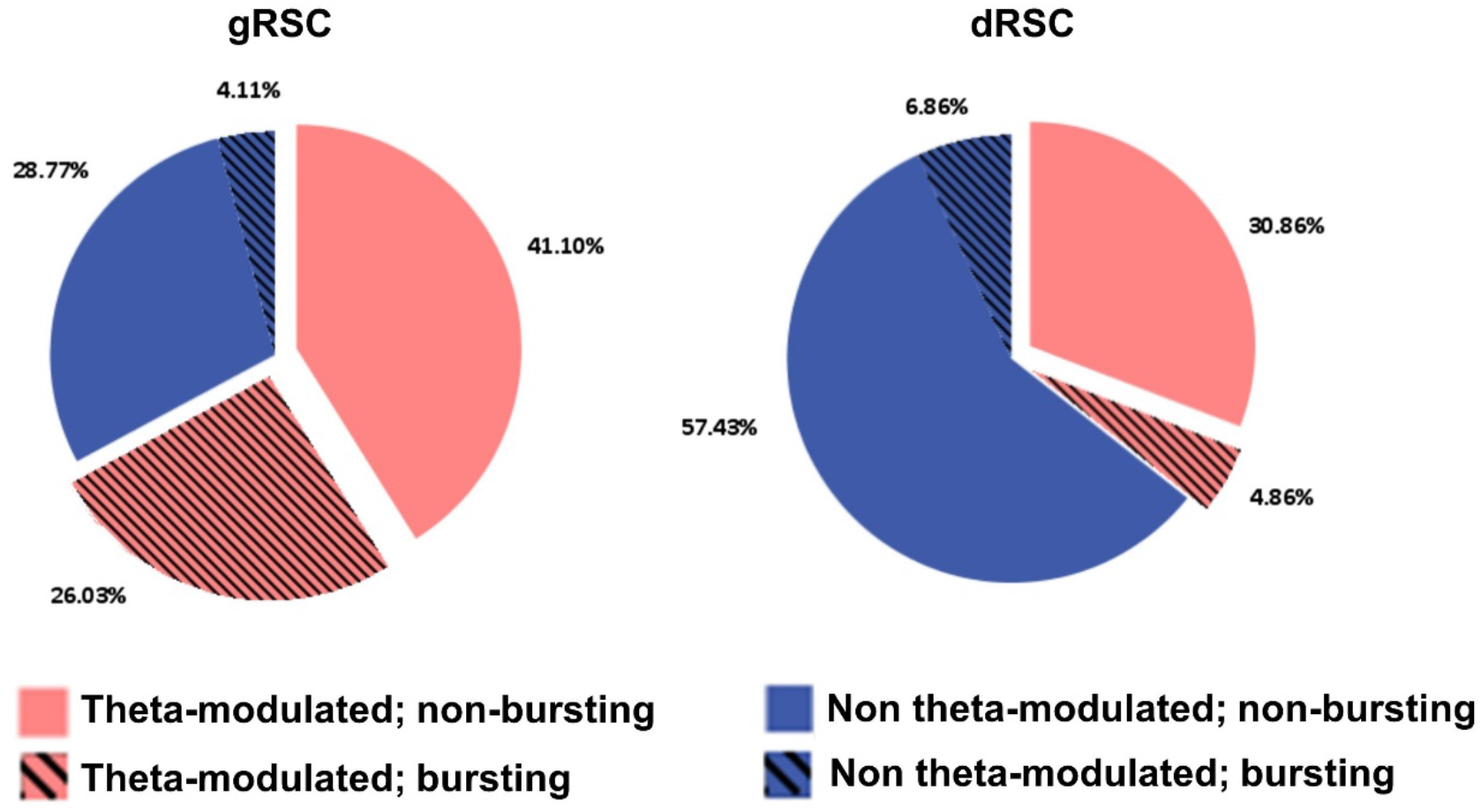
Theta-modulated cells are concentrated in the granular subregion of the RSC (gRSC). Theta-modulated and non-theta-modulated cells include bursting and non-bursting cells, divided based on peak ISI for the cell less than 10ms. gRSC: the majority of the cells fire coupled to the LFP theta, whereas most dysgranular RSC (dRSC) cells fire uncoupled from theta. gRSC also contains a higher proportion of bursting units, and the majority of these units are theta-bursting. In contrast, dRSC contains only a marginal subpopulation of bursting units, both theta and non-theta.

#### 3.1.5 Waveform differences: theta phase-locked cells have narrower waveforms

Finally, we examined whether theta-modulated and non-theta modulated cells in RSC were also morphologically distinct in their waveforms. We first examined the distribution of peak-to-trough widths by fitting a two-Gaussian component, and two conservative cutoffs were identified on this model to separate cells into narrow, broad and in-between (see Methods). We discarded 28 cases because spike waveforms did not reach time for repolarization. The analysis resulted in two modes (cell types): narrow spiking cells, 47.23%; broad spiking cells, 46.6%. A further 6.19% of cells were left unclassified based on values falling into the dip of the bimodal distribution (**Figure 7**). The calibrated Hartigan’s Dip Test discarded the null hypothesis of unimodality (DIP = 0.062; *p* < 0.001). We then compared the firing characteristics of cells in the two groups identified by the model. Cells classified as the narrow spiking type displayed a higher proportion of theta-modulated cells (narrow, 196/290; broad, 99/286; *χ*^2^ = 62.65, *p* < 0.001; **Figure 7**) and significantly stronger theta phase locking (IC: narrow, 0.14 ± 0.009; broad, 0.06 ± 0.003; z = -6.80; *p* < 0.001) compared to the broad type.

**Figure 7.**
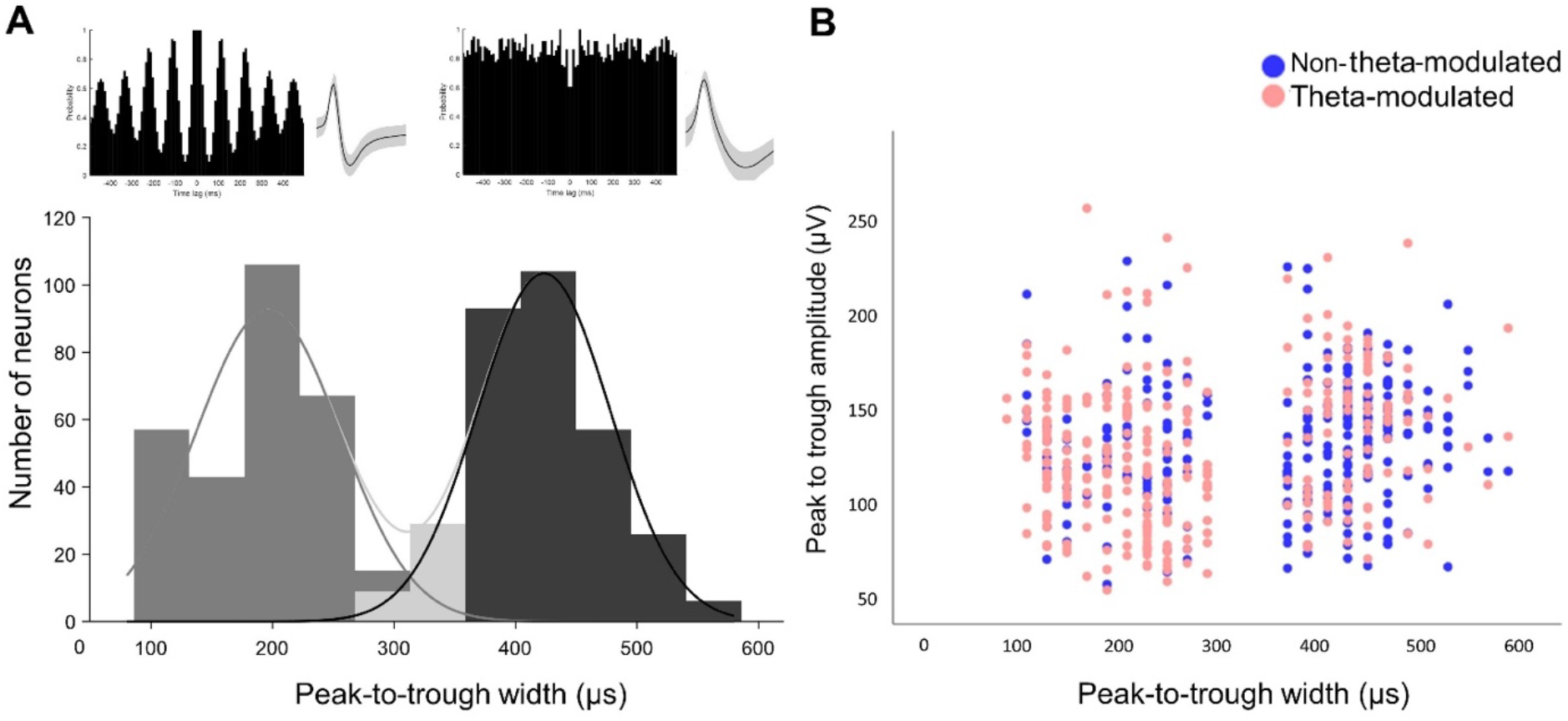
**Broad and narrow spiking cells display differences in theta-modulation strength**. **(A)** Distribution of the peak-to-trough width (µs) across all cells. Gaussian fits for the different modes (cell types) are displayed: broad (black), narrow (dark gray), and unclassified (light gray). Top, examples of autocorrelogram and averaged waveform for theta-modulated (left) and a non-theta-modulated (right) unit. Mean and standard deviation of the waveforms are shown. Theta-modulated cells tend to have spikes of shorter duration and a higher firing frequency than non-theta-modulated ones, suggestive of a higher prevalence of fast-spiking interneurons. **(B)** Waveform width of narrow and broad spiking cells in (A), plotted as a function of waveform amplitude (µV). A higher number of theta-modulated cells is found in the narrow group (n=196/290; 66.6%), whereas a higher number of non-theta-modulated cells is found in the broad group (n=187/286; 65.4%).

Cells were considered to be interneurons if the waveform peak-to-trough width was < 250µs and mean firing rate was > 5Hz (Henze et al., 2000; McCormick et al., 1985). We compared the proportion of interneurons between cell types and subregions. The proportion of interneurons was significantly higher among theta-modulated cells compared to non-theta modulated cells (theta-modulated: 63/321, 19.63%; non-rhythmic: 10/321, 3.12%; *χ*^2^ = 43.42, *p* < 0.001). As a result, this proportion was significantly higher in gRSC compared to dRSC, due to the anatomical organization of theta-modulated cells (gRSC: 39/292, 13.36%; dRSC: 12/350, 3.53%; *χ*^2^ = 21.45, *p* < 0.001). In agreement with the estimations based on comparisons of proportions, a two-way ANOVA conducted on waveform peak-to-trough widths found a main effect of cell type, *F*(1,1) = 20.12, *p* < 0.001, and brain region, *F*(1,1) = 18.38, *p* < 0.001, but no interaction, *F*(1,1) = 3.57, *p* = 0.06. According to these results, theta-modulated cells, predominantly found in gRSC, displayed considerably shorter waveform duration compared to non-theta modulated firing cells, predominantly found in dRSC (theta-modulated: 295.26 ± 9.05µs; non-theta-modulated: 363.99 *± 7.36*µs; WRS test, z = -6.85, *p* < 0.0001). Average peak-to-trough width for gRSC cells were as follows: theta-modulated (n = 196), 266.02 *±* 11.54µs; non-theta-modulated (n = 96), 343.54 ± 14.08µs. Average peak-to-trough width for dRSC cells were as follows: theta-modulated (n = 125), 341.12 ± 13.66µs; non-theta-modulated (n = 225), 372.71 ± 8.58µs. Between-subjects analysis confirmed a narrower average waveform in gRSC compared to dRSC (gRSC, 291.51 ± 9.26µs; dRSC, ± 7.39µs; WRS test, z = 6.68, *p* < 0.001). As seen in **Table 1**, this difference was repeated in the within-subject analysis, albeit in one animal it was not significant (R636; z = 1.45; *p* =0.15). Collectively, these results suggest that a higher number of theta-modulated cells are fast-spiking interneurons compared to cells that fire uncoupled from theta **(Figure 7)**. Theta-modulated cells are therefore in a position to influence firing rates of neurons located within the limits of the RSC, without transmitting rhythmic-modulation to the local head direction cell population, to which they might be connected.

### 3.2 Retrograde tracer experiment

We next compared, using retrograde tracing, the thalamic afferents to gRSC and dRSC to see if they differ, supporting that these are distinct circuits. The findings have been divided between those tracer injections approximately at the AP levels (-5.5 and -5.8 from bregma) of the electrodes used in the electrophysiological studies and those injections placed more rostrally or caudally within the RSC (see **Table 2**). Injections were performed in two different laboratories (UCL, n = 6 animals; Cardiff University, n = 17 animals), thus providing an internal replication for our findings (see **Supplementary Tables 3-4** for injection coordinates).

**Table 2.** List of all retrograde tracer injections. 23 animals with a total of 29 injections.

| *Injections at electrode level (n=13)* |  |  |  |  |
| --- | --- | --- | --- | --- |
| *Case No.* | *Injection site* | *Tracer* | *AV label* | *Laboratory* |
| 222#9 | dRSC s + d | CTB | AVVL | Aggleton; Cardiff |
| 222#10 | dRSC s + d | CTB | AVVL | Aggleton; Cardiff |
| 077941#13 | gRSC d | CTB | AVVL | Aggleton; Cardiff |
| 077942#14 | gRSC d | CTB | AVVL | Aggleton; Cardiff |
| 878 | dRSC, s + d | Ctb-488 | AVVL | Jeffery; London |
| 878 | gRSC, s (d) | Ctb-594 | AVVL/AVDM | Jeffery; London |
| 731 | dRSC, s + d | FG | AVVL | Jeffery; London |
| 731 | gRSC, s + d | Ctb-594 | AVVL/AVDM | Jeffery; London |
| 730 | dRSC, d | Ctb-594 | (AVVL) | Jeffery; London |
| 730 | gRSC s + d | FG | AVVL/AVDM | Jeffery; London |
| 717 | dRSC s + d; (M2) | FG | AVVL | Jeffery; London |
| 729 | dRSC s + d | FG | AVVL | Jeffery; London |
| 682 | gRSC s (d) | Ctb-594 | AVVL/AVDM | Jeffery; London |

| ***Injections at rostral and caudal locations (n=16)*** |  |  |  |  |
| --- | --- | --- | --- | --- |
| ***Case No.*** | ***Injection site*** | ***Tracer*** | ***AV label*** | ***Laboratory*** |
| 187#9 | gRSC d (s)<br>**(Rostral)** | BDA | AVVL | Aggleton; Cardiff |
| 64#3 | dRSC/M2/V2, s + d<br>**(Rostral + caudal)** | FB | AVVL | Aggleton; Cardiff |
| 172#27 | dRSC (M2/V2), s + d<br>**(Rostral + caudal)** | FB | AVVL | Aggleton; Cardiff |
| 172#28 | dRSC (M2/V2), s + d<br>**(Rostral + caudal)** | FB | AVVL | Aggleton; Cardiff |
| 225#4 | dRSC s + d<br>**(Rostral)** | CTB | AVVL | Aggleton; Cardiff |
| 225#12 | dRSC s + d/gRSC d<br>**(Rostral)** | CTB | AVVL | Aggleton; Cardiff |
| 223#1 | dRSC s + d<br>**(Rostral)** | CTB | AVVL | Aggleton; Cardiff |
| 223#4 | dRSC s+d/gRSC d<br>**(Rostral)** | CTB | AVVL | Aggleton; Cardiff |
| 227#22 | dRSC s+d<br>**(Rostral)** | FB | AVVL | Aggleton; Cardiff |
| 227#22 | dRSC d<br>**(Caudal)** | CTB | AVVL | Aggleton; Cardiff |
| 227#24 | dRSC s + d<br>**(Rostral)** | FB | AVVL | Aggleton; Cardiff |
| 227#24 | gRSC d<br>**(Caudal)** | CTB | AVVL | Aggleton; Cardiff |
| 227#16 | dRSC d<br>**(Rostral)** | CTB | AVVL | Aggleton; Cardiff |
| 227#16 | dRSC s+d/gRSC d<br>**(Caudal)** | FB | AVVL | Aggleton; Cardiff |
| 227#202 | gRSC (dRSC) s + d<br>**(Caudal)** | CTB | AVVL/AVDM | Aggleton; Cardiff |
| 227#204 | gRSC s + d<br>**(Caudal)** | CTB | AVVL/AVDM | Aggleton; Cardiff |
| FB, fast blue; CTB, cholera toxin subunit b; BDA, biotinylated dextran amine; FG, fluorogold. |  |  |  |  |

The injection site indicates the center of the tracer deposit; s and d indicate superficial and deep layers of the granular retrosplenial cortex (gRSC) or dysgranular RSC (dRSC). For deep injections, layer I was not involved. Cases 187#9, 64#3, 172#27, 172#28 had multiple injections at both rostral and caudal locations. Those regions and layers in parenthesis indicate weak involvement in the injection site. The third column indicates the subfield of the anteroventral thalamic nucleus (AV) with densest cell label resulting from the injection. AVDM, dorsomedial subfield of the AV; AVVL, ventrolateral subfield of the AV.

#### 3.2.1 Injections at the level of recording

Thirteen retrograde tracer injections (n=10 animals) targeted RSC at approximately the same AP levels as the recording sites (for representative injections see **Figure 9**; all injection sites reported in **Table 2**). In three cases (730, 731 and 878), dual injections were made, one in dRSC (n = 6) and the other in gRSC (n = 7; **Table 2**). For dRSC, all injections included portions of both deep and superficial layers except a single injection (case R730) restricted to the deep layers.

In all cases, retrogradely labelled cells were present in the ipsilateral thalamus, with varying amounts of label in AD, AV, and the anteromedial nucleus (AM). Whereas both gRSC and dRSC injections consistently resulted in widespread AD labelling, the distribution of labelled cells within AV differed between the two injection sites. This difference within AV is the focus of the results.

In all cases with injections largely confined to dRSC, retrograde cell label was virtually restricted to the ventrolateral (AVVL) subfield of the AV nucleus (e.g., cases 222#10, 222#9). In contrast, in four of the six cases that targeted gRSC (cases 878, 731, 730, 682), retrograde cell label was observed in both AV subfields, i.e., AVVL and dorsomedial AV (AVDM). The two remaining gRSC cases (077941#13, 077942#14) contained label in both AVVL and AVDM, but the preponderance was in AVVL (**Figure 9**).

The overall pattern of differences (AVVL to dRSC; AVVL and AVDM to gRSC) can be observed in **Figure 9**, which provides a schematic depicting the AV area with the densest cell body labelling across the various injection cases (label in AD is not shown). In further support of this difference in AV innervation, **Figure 10** shows photomicrographs of the two different patterns of AV labelling in a single case (878) that had injections in both dRSC and gRSC.

**Figure 8:**
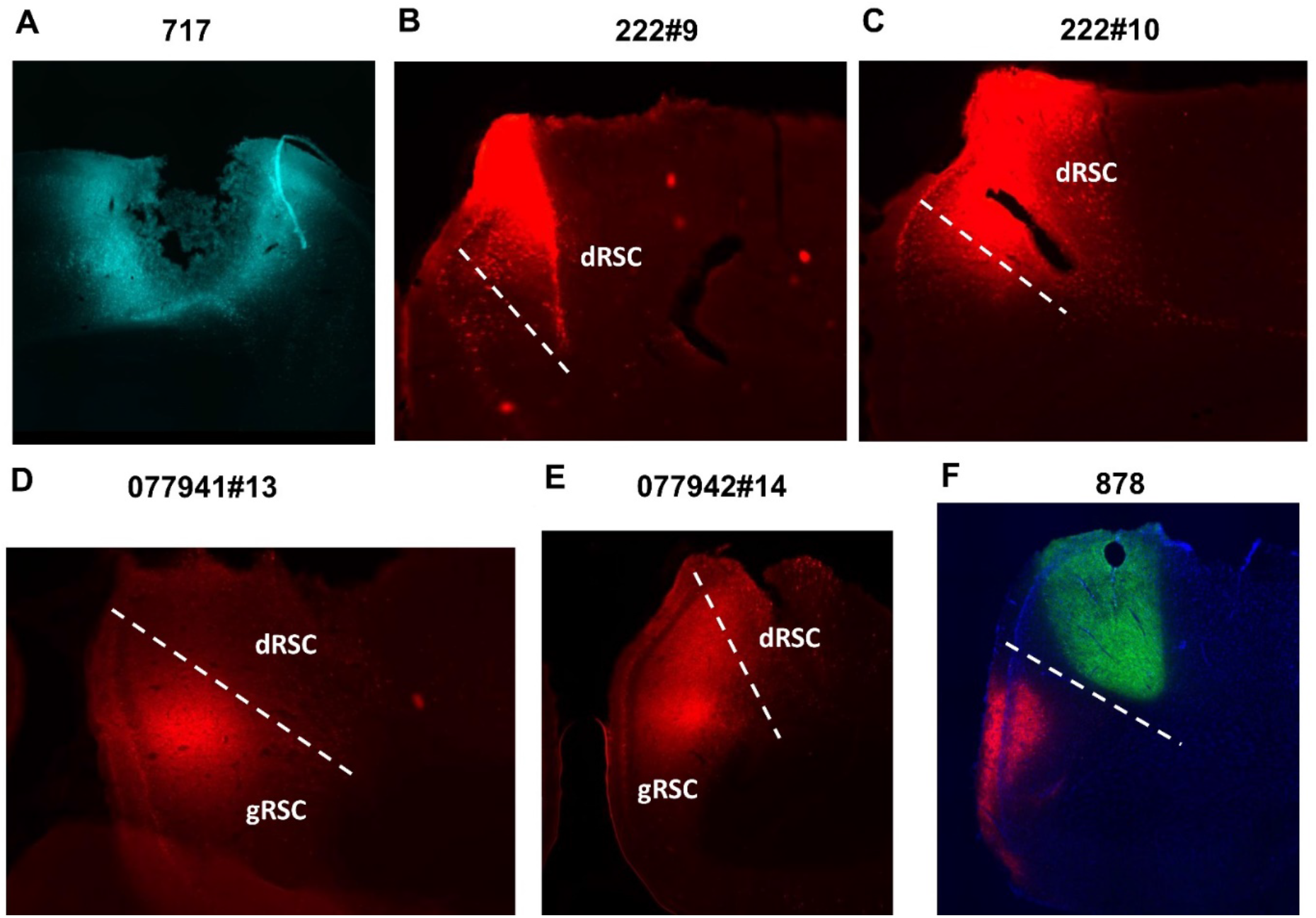
Injections in retrosplenial cortex (RSC) were made at the electrode recording levels. Cholera toxin subunit b (CTB) and fluorogold (FG) injections in **(A-B-C)** dysgranular RSC (dRSC, both superficial and deep layers) and **(D-E)** granular RSC (gRSC, mainly deep layers). **(F)** Dual injection in granular (mainly superficial layers) and dysgranular (superficial and deep layers) RSC. AP level: -5.5mm.

**Figure 9.**
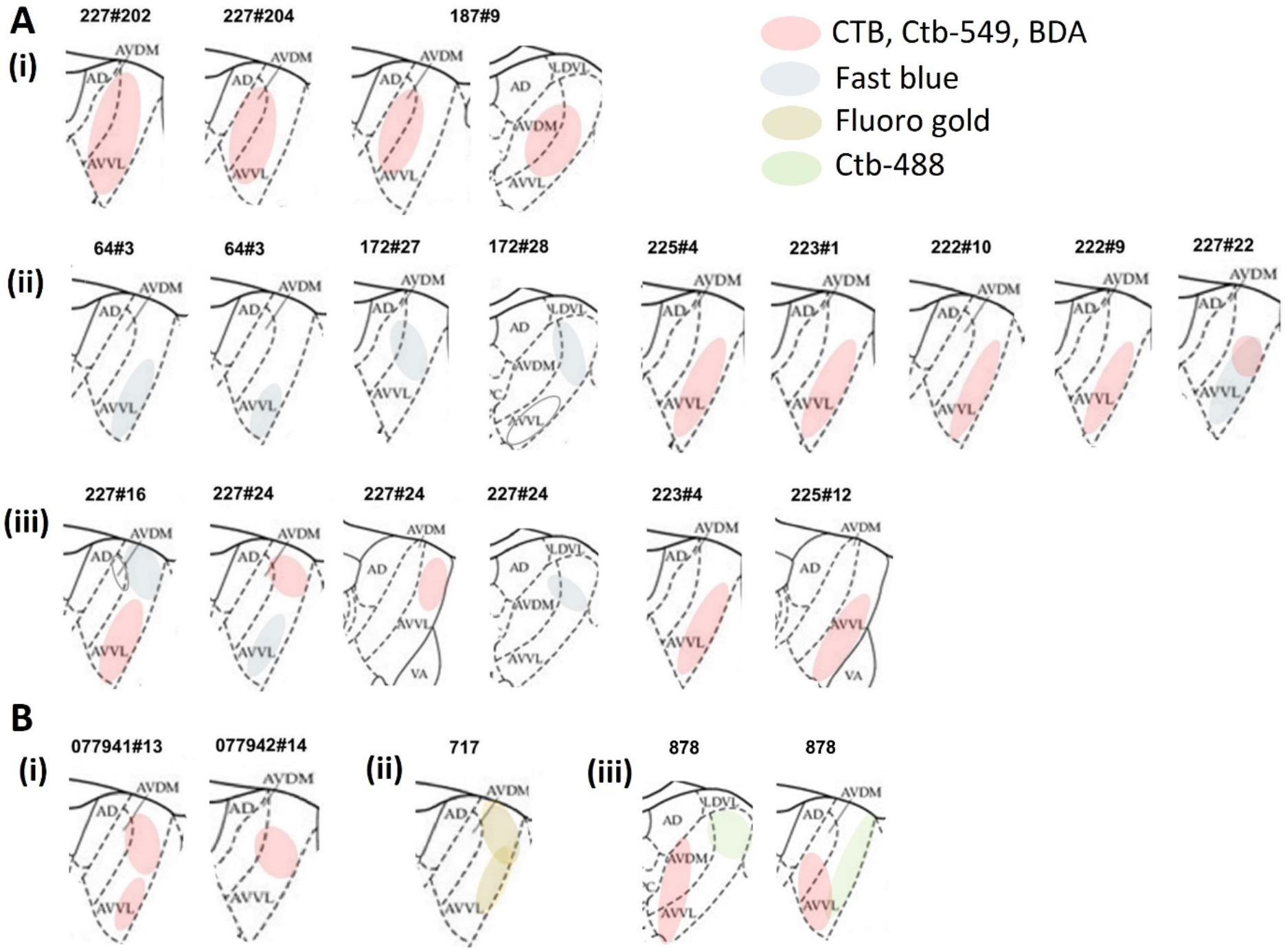
Ventrolateral AV (AVVL) widely projects to both gRSC and dRSC, dorsomedial AV (AVDM) projections are more restricted to superficial layers of gRSC. Schematics showing areas of retrograde cell labelling in AV**. (A)** Tracer injections rostral or caudal of electrode level; **(B)** injections at the electrode level. **(i)** AV label in cases of single tracer injection in gRSC. **(ii)** AV label in cases of single injection or multiple injections (127#27, 127#28, 54#3, 227#22) in dRSC. **(iii)** AV label in cases with combined gRSC and dRSC tracer injections. Cases 227#16, #24 and 878 have dual injections placed in dRSC and gRSC. Cases 223#4 and 225#12 have a combined injection in both gRSC and dRSC. Areas of label are color coded according to tracer type (see legend). For cases 227#16 and 172#28 there was weak additional labelling in AVDM and AVVL respectively, resulting from FB injection in dRSC (empty circles). Abbreviations: AD, anterodorsal thalamic nucleus; AVVL and AVDM, ventrolateral and dorsomedial subfields of the anteroventral thalamic nucleus; VA, ventral anterior thalamic nucleus; LDVL, ventrolateral subfield of the laterodorsal thalamic nucleus.

**Figure 10.**
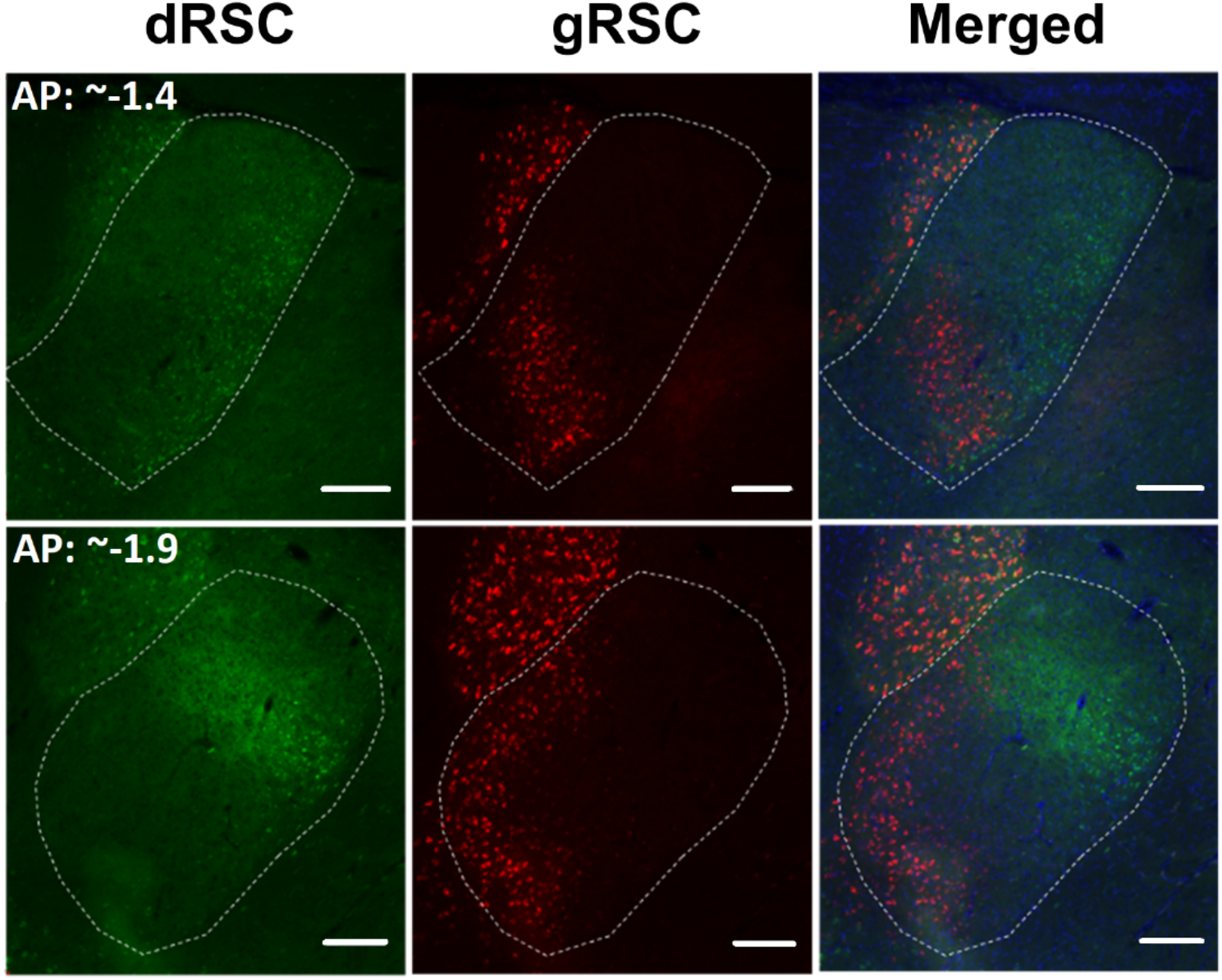
Cell label is seen in AVDM following injections in superficial layers of gRSC, and in AVVL following dRSC injections. Dual retrograde tracer injections into dRSC (CTB-488 green) and gRSC (CTB-549 red), with AV borders indicated. See injection sites in Figure 8 (Case 878). The photomicrographs show separate and combined images for two different AP levels within the same animal. Cells labelled with CTB-488 (green; dRSC) are concentrated in lateral AV, corresponding to AVVL, whereas CTB-549 label (red, gRSC) is also present more medial, i.e., in both the AVVL and the AVDM subfields (see Figure 9 for schematic). Scale bars 200µm.

We then correlated the variations in labelling with variations in injection site. For gRSC, in the four cases with combined AVDM/AVVL label the injection site included both deep and superficial layers (cases 878, 731, 730, 682), whereas in two cases where the label was predominantly in AVVL, the gRSC injections were restricted to the deep layers (cases 077941#13, 077942#14; see **Figures 8 D, E; 9**).

#### 3.2.2 Injections at more rostral and caudal RSC locations

In a further sixteen retrograde tracer injections (n=13 animals), we targeted portions of RSC >0.5 mm rostral or caudal to the electrophysiology recording sites (**Table 2;** for representative injections see **Supplementary Figures 2-3**). Of these, six injections were from three cases with dual tracer injections at different AP levels (**Supplementary Figure 2**). Nine tracer injections seemed restricted to the dRSC, four to the gRSC and three included both RSC subregions (**Supplementary Figure 3**). These dRSC injections included both superficial and deep layers, except for two cases where the injections appeared restricted to deep dRSC (rostral 227#16, caudal 227#22). Irrespective of AP level, we observed consistent labelling differences after dRSC and gRSC injections (**Supplementary Figure 2**) that closely matched those differences seen when the injections were placed at the level of the electrodes. Once again, dRSC injections were consistently associated with AV label in AVVL (only occasionally extending marginally into AVDM towards its dorsal border), while gRSC injections resulted in label in both AVVL and AVDM **(Figure 9)**.

Further examples of these differential patterns of AV label can be see in **Figure 11**. This figure contrasts when a tracer injection involved both dRSC and gRSC, versus when it was located in just the dRSC subregion. An injection of CTB involving both RSC subregions resulted in extensive cell labelling across both AVDM and AVVL (case 227#202; **Figure 11 A, B**). Meanwhile in a case with dual injections (case 227#16), the rostral dRSC injection (CTB, red) resulted in ventral AVVL label, while the caudal fast blue injection (primarily dRSC but also deep gRSC) labelled dorsal AVVL (case 227#16; **Figure 11 C, D**).

**Figure 11.**
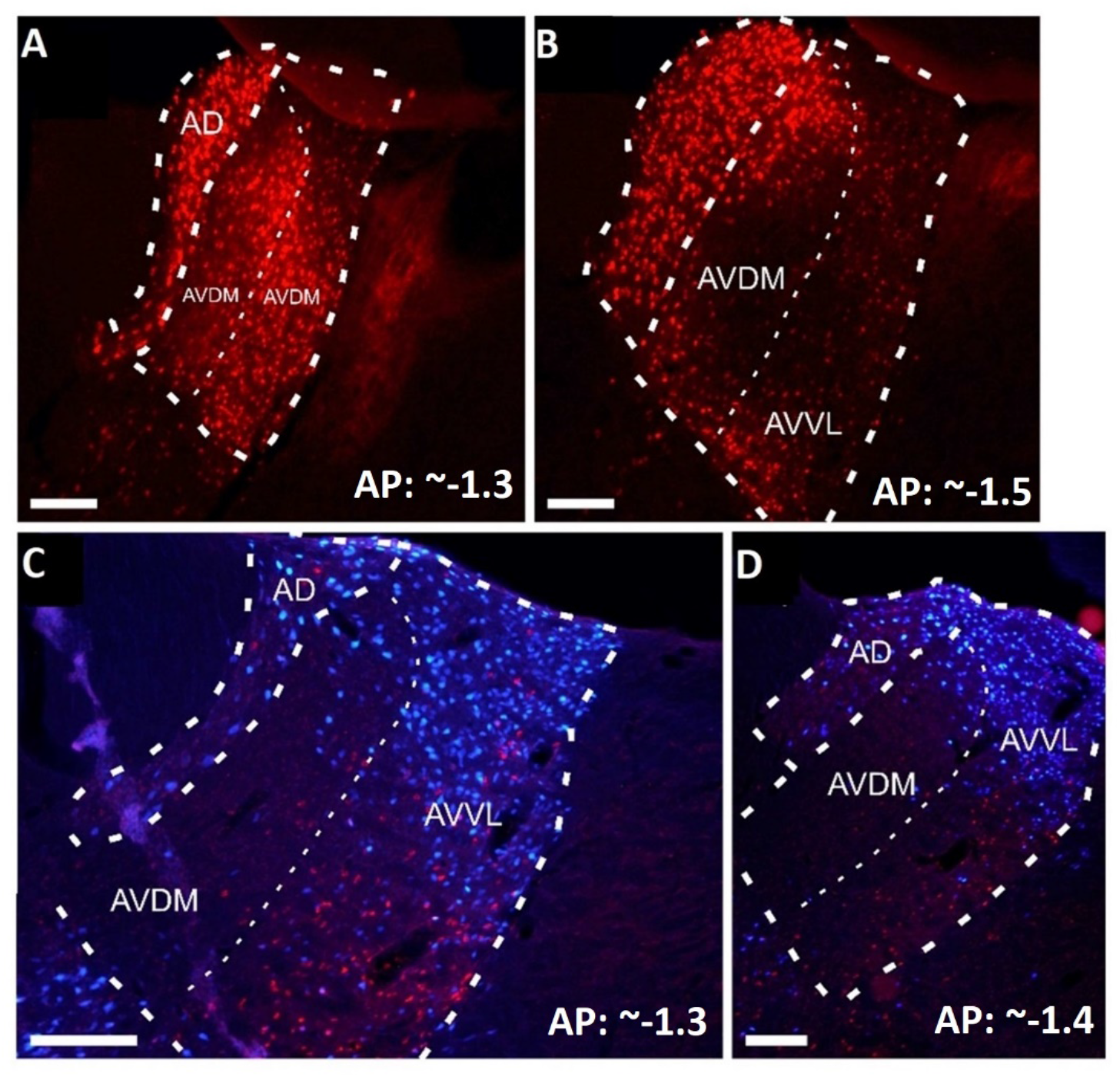
Injections into both deep and superficial layers of gRSC (upper) result in more extensive and combined AVDM/AVVL label than is seen after dRSC injections (lower). AVDM Photomicrographs depicting retrograde cell label after retrosplenial injections. **(A-B)** Cell label following a CTB injection in gRSC (case 227#202, superficial and deep layers). Retrograde labelled cells are seen in both AV subfields. See injection sites in Supplementary Figure 3. **(C-D)** Retrograde labelled cells following dual injections in dRSC at anterior (CTB) and posterior (FB) portions (case 227#16). See injection sites in Supplementary Figure 2. For both injections, cell labelling is predominantly present in the AVVL subfield, with more scattered cell label in the AVDM subfield (see Figure 8 for schematic). Two different APs are shown for the same animal. AD, anterodorsal thalamic nucleus; AVDM, dorsomedial subfield of AV; AVVL, ventrolateral subfield of AV. Scale bars 200µm.

In case 227#24 (**Figure 12**) separate injections were placed in rostral dRSC (fast blue, involving deep and superficial layers) and in caudal gRSC (CTB, deep layers; see also schematic **Figure 9**). This and other cases with dual injections positioned at rostral and caudal portions of the RSC (cases 227#22 and 227#24, 227#16; **Figures 11-12**) also supported a previously described topographical pattern (Shibata, 1993): namely, that ventral AV preferentially projects to the rostral RSC whereas dorsal AV targets caudal RSC (see **Figure 11 C, D**). This same gRSC injection (227#24) showed a concentration of AVVL cell label (**Figure 12**), whereas three other cases with gRSC injections resulted in combined AVVL/AVDM cell label (cases 227#204, 227#202,187#9; as also described above for the gRSC injections close to the AP of the recording sites see **Figures 8-9**).

**Figure 12.**
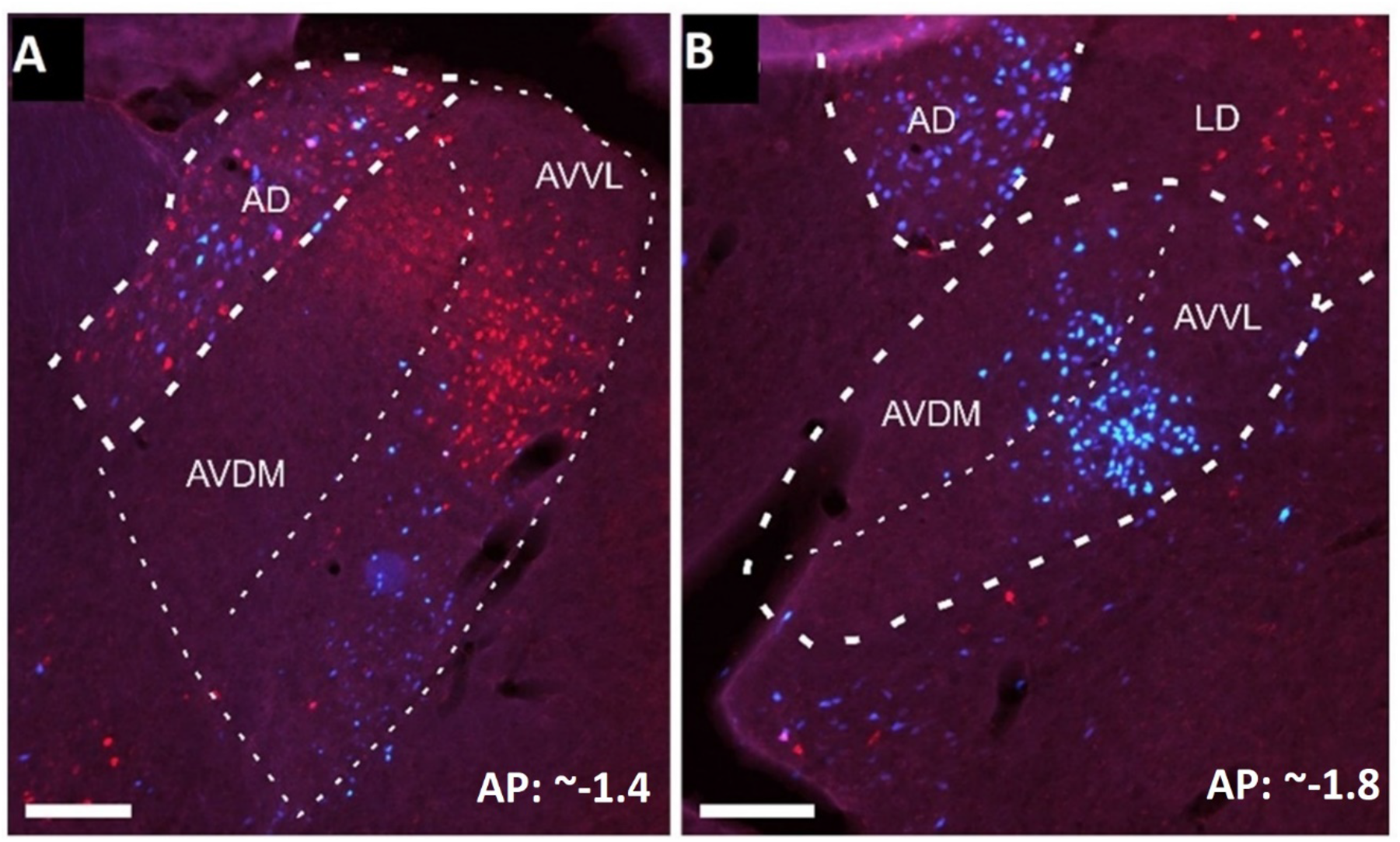
Injections positioned in deep layers of gRSC only and more moderately label a cell cluster in the AVVL, similarly to dRSC injections. Photomicrographs of AV label for case 227#2, after dual injections in gRSC (deep) and dRSC (superficial and deep), at two different AP levels. See injection sites in Supplementary Figure 2. Fast blue (blue) label originated from an injection centred in rostral dRSC (both superficial and deep layers) whereas cholera toxin-b label (red) originated from an injection centered in caudal gRSC (deep layers only). The majority of fast blue cell label is in AVVL subfield (see Figure 9 for schematic). Unusually for a gRSC injection, the retrograde label (red) was also largely confined to AVVL, potentially reflecting the deep location of the tracer injection. Two different APs are shown for the same animal. Abbreviations: AD, anterodorsal thalamic nucleus; AVDM, dorsomedial subfield of AV; AVVL, ventrolateral subfield of AV; LD, laterodorsal thalamic nucleus. Scale bars 200µm.

Further inspection of these cases revealed that the single case with restricted AVVL label (227#24) had a gRSC injection largely confined to its deep layers, whereas the three cases with further AVDM involvement had either obvious (case 227#204; 227#202) or likely (case 187#9 – see **Supplementary Figure 3**) involvement of superficial layers in the injection sites. These observations support the earlier conclusion that it is particularly the superficial layers of gRSC that receive AVDM projections. In further agreement, the three remaining cases with combined gRSC/dRSC injections had cell label in AVVL (but not AVDM) and in these cases the gRSC portion of the injection sites seemed largely contained within the deep layers (cases 225#12, 225#4, 227#16).

## 4.0 Discussion

As part of an ongoing attempt to understand the contribution of retrosplenial cortex to spatial memory processing, we investigated what distinguishes its two principal subregions, granular (gRSC) and dysgranular (dRSC). Recent findings of landmark-sensitive directional neurons restricted to the dysgranular region only (Jacob et al., 2017) have suggested that RSC may comprise two subcircuits, one (gRSC) more connected to hippocampus and the other (dRSC) more connected to sensory regions. We thus investigated whether there are electrophysiological or anatomical differences that shed light on this dichotomy. Electrophysiologically, we compared the temporal firing characteristics, waveform characteristics and relation to the local LFP of non-directional cells recorded from gRSC and dRSC during free exploration in rats, finding greater coupling with theta oscillations in gRSC compared to dRSC. Anatomically, we used retrograde tracing to label gRSC and dRSC projections from one of the main inputs to RSC, the anterior thalamus, to see if there might be differential projections from this region; we found that gRSC was distinguished by a projection from the dorsomedial subfield of the anteroventral thalamus (AVDM) that was much weaker or missing following the dRSC injections. Below, we examine these findings, and then conclude with some speculations about the possible functional role of these two thalamocortical subcircuits.

### 4.1 Increased theta coupling in gRSC

Comparison of the two subregions (in both pooled and single subject analysis) revealed that gRSC had a much higher mean IC (index of theta phase-coupling), firing frequency and spike-bursting propensity, and theta-modulated cells in this region exhibited considerably larger theta-modulation depth. However, we found no difference in local field potential (LFP) theta oscillations between the two subregions, highlighting a dissociation between theta rhythms in the LFP and oscillations present at the single cell level (i.e., theta rhythms are not generated uniquely from local ensemble activity within gRSC, in line with results of Young and McNaughton, 2009; Borst et al., 1987, Talk et al., 2004). These observations raise two questions: what causes these differences and what, if any, are their functional implications?

One possibility is that these signals are a consequence of differential network connectivity between gRSC/dRSC and the medial septal theta “clock.” Septal GABAergic neurons express theta-modulated firing and innervate both CA1 and gRSC – but dRSC much less so – with a common projection pattern (Unal et al., 2015). They selectively target and inhibit GABAergic interneurons, resulting in a rhythmic disinhibition of the principal neurons, the activity of which thus becomes phase-locked to septo-hippocampal theta. Rhythmic disinhibition supplied by the medial septum has been demonstrated to drive hippocampal theta activity (Hangya et al., 2009) and could offer a similar explanation for the concentration of theta cells in gRSC. Given the lack of septal input to dRSC, some dRSC neurons might become locked to theta indirectly via cell-to-cell projections from gRSC theta cells.

An alternative possibility is that theta is modulated by the strong anterior thalamic projection to RSC (both gRSC and, more moderately to, dRSC). Like cells in the medial septum, theta rhythmically discharging cells in the AV fire locked to the local and hippocampal theta oscillations, and these oscillations are strongly coherent (Albo et al., 2003; Vertes et al., 2001; Tsanov et al., 2011, 2010). The medial mammillary bodies distribute massively to the AV via the mammillothalamic tract (Seki and Zyo, 1984; Shibata 1992), and projects to the AV a theta signal that combines tegmental, septal, and hippocampal theta activity (see reviews by Vertes et al., 2004; Vertes and Kocsis, 1997). A role of the mammillary bodies-AV axis in complementing the septo-hippocampal theta system (Zakowski et al., 2017; Ruan et al., 2017) may explain the occasional report of theta oscillations in gRSC that persist or are enhanced in the absence of hippocampal theta (i.e., during hippocampal large amplitude irregular activity, Young and McNaughton, 2009; or electrolytic septal lesions, Borst et al., 1987). A third possibility is that gRSC inherits theta rhythmicity via its stronger afferent connections with the dorsal hippocampus, both from CA1 (Haugland et al., 2019; van Groen and Wyss, 1990; Opalka et al., 2020; Unal et al., 2015) and subiculum, which is the major output of the hippocampus and sends a strong projection to gRSC, but not dRSC (Meibach and Siegel, 1977). We emphasize that these options are not mutually exclusive. Thus, the AV may exert an excitatory rhythmical influence on target cortical neurons receiving converging hippocampal or septal projections (Yamawaki et al., 2019), which may be crucial for allowing a subcortically-mediated regulation of coordinated hippocampus-RSC communication (Chambers et al., 2021).

The second major question arising from the differential theta coupling observed between gRSC and dRSC is what, if any, might be the functional consequences. Theta is widely thought to have the function of synchronizing the excitability of brain regions throughout the hippocampal formation (Korotkova et al., 2018), perhaps to facilitate information transfer or synaptic plasticity. Within hippocampus itself, each theta cycle contains a sequential activation of hippocampal neurons (usually manifesting as place cells) and may function to organize sequential information as part of a spatio-temporal memory system. It may be that gRSC with its greater hippocampal connectivity has the role of mediating between incoming sensory information and ongoing spatial computations, with theta facilitating the transfer and maintaining the spatio-temporal organization. We will return to this point later.

### 4.2 Differential AV projections to gRSC and dRSC

Our dataset of retrograde tracer injections adds to previous anatomical descriptions of anterior thalamic inputs to RSC. Overall, in agreement with previous studies (Shibata, 1993), a topography emerged from our tracing studies such that ventral AV projects to rostral RSC while dorsal AV projects to caudal RSC. Within the AV projections, we found a clear differentiation of AVDM and AVVL projections into the gRSC and dRSC. The AVVL subfield targeted both retrosplenial subregions, providing moderate fiber input to both superficial and deep layers (see also Van Groen and Wyss, 1990a, 1992, 2003; Shibata, 1993). In contrast, the AVDM subfield provided a selective projection to the gRSC; mainly to the superficial layers (Shibata, 1993). Overall, the present results argue for the presence of two related but distinct AV-RSC subcircuits: AVVL to dRSC, AVVL and AVDM to gRSC.

Two anatomical facts are relevant for clarifying the observed variations in the electrophysiology data between gRSC and dRSC. First, the difference in the overall projection strength to the two RSC subregions should be noted, in addition to the topographical organization of AV projections from within AV to each of these subregions (Shibata et al., 1993). In relation to the first point, our data demonstrate stronger AV projections to gRSC compared to dRSC (see also Van Groen and Wyss, 2003). Second, there are differences in the source of these projections from different AV subfields. Superficial gRSC layers receive a distinct projection from AVDM, which sends no or only moderate projections to dRSC. In contrast, dRSC only receives widespread AVVL projections. Hence, observed differences in theta modulation across the RSC might not simply be an effect of variations in AV fiber density (stronger AV to gRSC) but may also be explained by considering anatomically distinct cell populations within the AV projecting to the two RSC subregions.

### 4.3 Two interacting thalamocortical subcircuits

Previous behavioral, connectivity and neuroanatomical evidence, together with our current findings, combine to indicate that the RSC is not a unitary structure (see Aggleton et al., this issue, for review). gRSC is more strongly coupled with the hippocampal formation, as evidenced by both anatomical connectivity and theta modulation: the latter of which accords with the fact that theta is a distinguishing feature of neurons in brain regions that are linked to the hippocampal spatial system (Burgess et al., 2002; Buzsáki, 2002, 2005, Hasselmo and Eichenbaum, 2005; Buzsáki and Moser, 2013). Our findings add to this picture by showing that there is a selective contribution of the AVDM to this network, via gRSC. Future work will need to determine the distinct physiological properties of AVDM neurons, and the consequence on gRSC processing of removing this projection.

By contrast, dRSC is dominated by visual interactions (Fischer et al., 2020; Powell et al., 2020; Keshavarzi et al., 2021, van Groen and Wyss, 1992, 1990b,c; Van Groen and Wyss, 2003; Cooper and Mizumori, 2001; Pothuizen et al., 2009). This conclusion is bolstered by the recent findings of Jacob et al. (2017), who demonstrated stronger visual responsiveness in dRSC directional signals, which suggested a partial uncoupling from the main, hippocampally interacting head direction signal. The question thus arises: why does the brain maintain these separable subsystems?

One possibility is that this separation allows for the comparison of two information streams that may agree or differ in their content, and thus allow for comparison and appropriate error correction. The streams in RSC comprise interoceptive signals arising from subcortical regions and exteroceptive signals coming in from the sensory periphery. A specific example of these converging information streams is provided by the RSC head direction signal, which arises from convergence of self-motion inputs (primarily interoceptive cues such as vestibular signals, motor efference and proprioception) and static environmental information from the sensory cortices (Yoder and Taube, 2014). The findings of Jacob et al (2017) that directionally tuned neurons in dRSC operate independently of this “consensus” head direction signal have suggested that RSC may be the site of weighted cue integration between the current estimate of head direction and incoming sensory information (Page and Jeffery, 2018). A similar principle may account for RSC processing more generally: that is, the gRSC input from thalamus may supply an estimate of a current state, the dRSC supplies current sensory input, and intrinsic connections between the subregions could allow for integration and weighted cue combination. Thus, via intrinsic granular-dysgranular interconnections (Shibata et al., 2009; Sugar et al., 2011), RSC can mediate exchange and integration of information between more cognitive and more perceptual computations.

### 4.4 Conclusion and future directions

Our results detail differences in theta-related spiking between gRSC and dRSC along with the existence of two interdependent AV-RSC subnetworks, suggesting two distinct streams of information to the RSC. We propose that these conform to a more interoceptive stream that conveys the results of internal processing and a more exteroceptive stream that conveys the state of the external world as assessed by the senses. The cognitive stream is characterized by close connectivity with the hippocampal system, as well as a unique input from the AVDM. This convergence may represent a key aspect of RSC function, consistent with evidence from human neuroimaging studies showing a role for RSC in monitoring landmark stability (Auger et al., 2012) and integrating across scenes and reference frames (Sherrill et al., 2013; Marchette et al., 2014; Bicanski and Burgess, 2016; Byrne et al., 2007). In the future, a complete understanding of the limbic system will necessarily need to consider the inter-related functions of these subnetworks, rather than the function of either AV or RSC as a whole.

## Acknowledgments

The authors would like to thank Roddy M. Grieves and James Street for the continued advice throughout these experiments and help with programming. We thank Pierre-Yves Jacob for his contribution to the collection of electrophysiology. The authors are grateful to Brook Perry for advice on the LFP analysis.

This work was supported by three Advanced Investigator Grants: KJ was supported by grants from Wellcome (103896AIA) and the Royal Society (IEC\R2\181140). ASM was supported by a Wellcome Trust Senior Research Fellowship (WT 110157/Z/15/Z). JPA and MM were supported by grants from Wellcome (103722/Z14/Z) and the UK BBSRC (BB/T007249/1). EL was supported by a Medical Sciences Graduate School Studentship, University of Oxford: Clarendon Fund – Department of Experimental Psychology –Somerville College Mary Somerville Graduate Studentship (SFF1718_CB2_ MSD_1074636). N.Z. was supported by a scholarship from the China Scholarship Council (201608000007).

## Data sharing

Data supporting the findings of this study are available on request from the corresponding author, Eleonora Lomi. Email address

## Authors contributions

All authors discussed and reviewed the results. E.L. wrote the manuscript with input from all authors, who reviewed and edited the manuscript. In addition:

EL: Conceptualization and investigation of the project; Analysis of electrophysiology data; Data interpretation; Visualization.

MM: Collection and analysis of neurotracing data; Data interpretation.

HYC: Collection and analysis of neurotracing data.

NZ: Collections of electrophysiology data.

JPA: Funding acquisition; Resources; Supervision.

ASM: Conceptualization and administration of the project; Funding acquisition; Resources; Supervision.

KJ: Conceptualization and administration of the project; Protocol design; Funding acquisition; Resources; Supervision.

