## Supplementary Materials for "Evidence for two distinct thalamocortical circuits in retrosplenial cortex"

### 1.0 Electrophysiology recording experiment

#### **1.1 Histological considerations**

The AP level of the tetrodes for each rat (n=4) was inferred from the shape of the fibers of the cingulum (Cg), by referencing to the atlas of Paxinos and Watson (2007). Although the shape of the hippocampus would indicate more anterior APs for R887 and R911 (i.e., AP of -4.0 mm from bregma), the AP level of the actual tract in the cortex above the hippocampus is more posterior, as the caudal end of Cg disappears at a coronal AP level of -5.5mm. Moreover, for both rats, the two hemispheres are separated by the deep central sulcus, suggesting that recordings in the cortex could not have been taken from APs < -5.5mm (Figure 1). Hence, given that it is possible that the brain was cut at an angle, we did not use the hippocampus shape as a guide to determine the AP.

For two animals (R602, R911) the electrodes mostly sampled superficial gRSC layers (I-IV), although we cannot exclude potential involvement of other layers. For the remaining two animals (R636, R887), sampling largely included both superficial and deep gRSC layers. These two sets of animals may also have comprised different recording sites within gRSC: while both exhibited sampling of dRSC and gRSC area b, additional gRSC area a sampling may have occurred only for the two animals with deeper implants (see main text **Figure 1**).

#### **1.2 Phase preference analysis**

Within each animal (single subject analysis), theta-rhythmic cells (see Methods, IC > 99^th^ percentile of shuffle distribution) in gRSC exhibited a consistent phase preference, albeit the exact preferred phase is hard to determine because theta was not measured from a fixed reference point across animals, due to different placements of the electrodes. In two animals (R636, R887), LFP-spike coupling occurred on the descending phase just before the trough of the rhythm, with preferred theta phases at 143.49*^◦^* and 162.04*^◦^*, and significant R-vectors of 0.75 and 0.34 (Rayleigh test, R636 (n = 79): z = 9.52; R887 (n = 10): z= 5.74; all *p* < 0.01). In the remaining two animals (R602, R911), LFP-spike coupling occurred on the ascending phase just before the peak of the rhythm, with means at 308.69*^◦^* and 343.70*^◦^*, and significant R-vectors of 0.31 and 0.58 (Rayleigh test, R602 (n= 41): z = 3.85, *p* = 0.02; R911 (n = 66): z = 22.1, *p* < 0.0001). Differences in preferred spiking phases between these two sets of animals are illustrated by the circular distributions (**Supplementary** **Figure 1).** Even by eye, it is clearly apparent that preferred theta phases of theta-rhythmic cells fell in the same 180*^◦^* hemisphere for two animals (n = 89; WW test, F = 4.38, *p* > 0.05), while they fell in the 0/360*^◦^* hemisphere for the other two animals (n = 107; F = 0.72. *p* > 0.05). The preferred hemisphere is highlighted on each circle. A WW test confirmed that the mean phase of the two sets of animals differed significantly from each other, suggesting firing out of phase (F = 164.94, *p* < 0.001). The dRSC theta-rhythmic cells displayed highly variable phase preference in two animals, indicated by uniform phase distributions (Rayleigh test, R602 (n = 86): z = 0.03; R636 (n = 7): z = 1.15; all *p* > 0.05). Only in two cases (R887, R911), dRSC theta-rhythmic cells fired consistently on the same theta phase, with means at 63.41*^◦^* and 295.15*^◦^*, and significant R-vectors of 0.63 and 0.89, respectively (Rayleigh test, R887 (n = 13): z = 5.17; R911 (n = 19): z = 15.2, all *p* < 0.01). Note that for these two animals, the firing of theta cells converged on the same phase (theta troughs for R887; theta peaks for R911) for gRSC and dRSC recordings (**Supplementary** **Figure 1**).

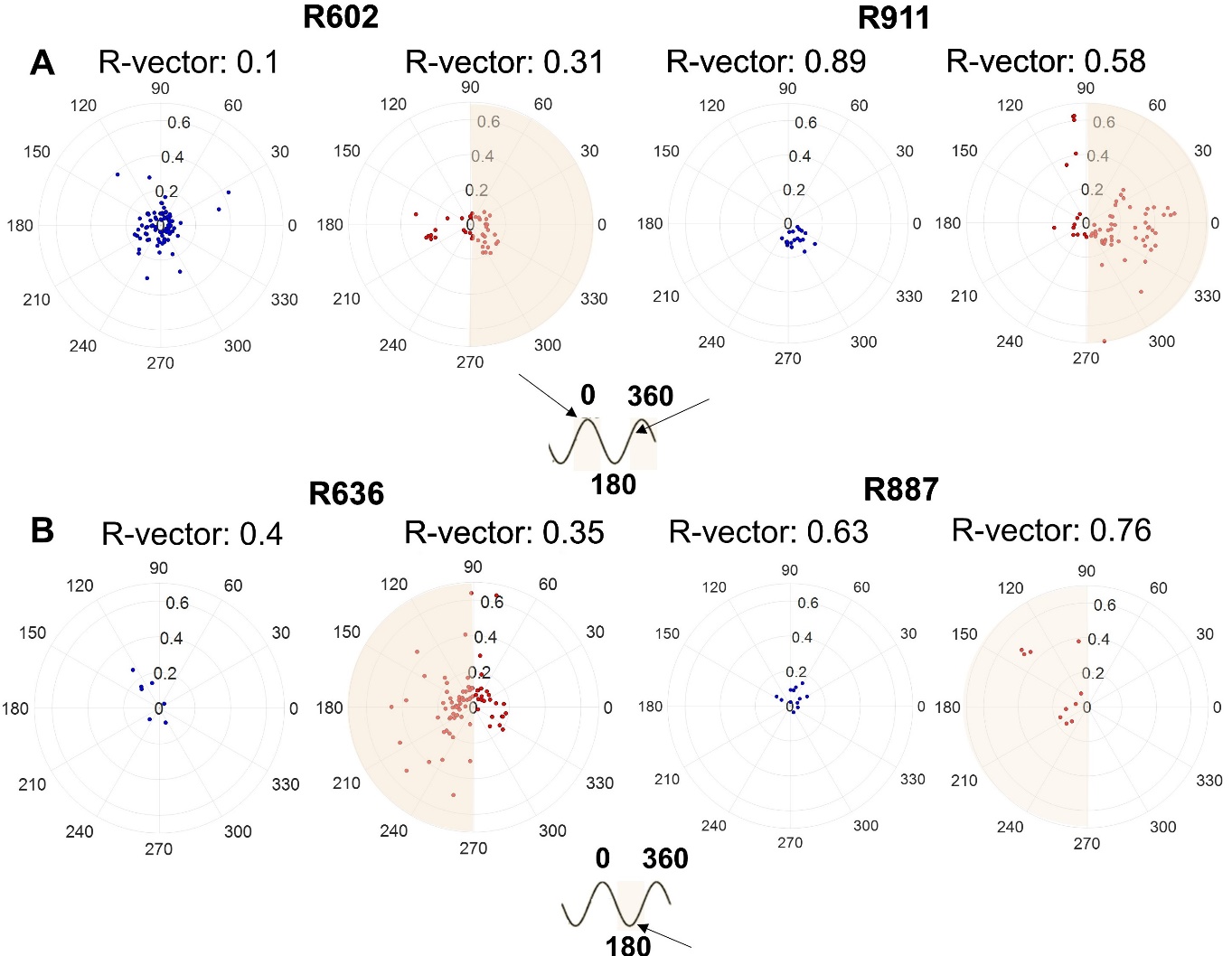

**Supplementary** **Figure 1. Distribution of preferred phases of single neurons.** For each animal, gRSC is shown to the right; dRSC to the left. Rat number at the top. Each marker is the preferred theta phase of a cell, and the radius of each marker is its theta-modulation depth (measured by the index of theta phase-coupling, IC). For each animal, the preferred hemisphere is highlighted in tan. gRSC: in addition to the stronger theta modulation within each cell (markers falling further away from the centre), across cells there was a more consistent phase preference (unimodal phase distributions within each animal). Distributions were unimodal for all animals (Rayleigh tests, all *p* < 0.01). Across all gRSC cells, preferred theta phases converged in the 0*^◦^* region for two animals **(A)** and in the 180*^◦^* region for the other two animals **(B)**, suggesting firing on opposite peaks of the cycle between animals. dRSC: cells displayed weaker theta modulation within each cell (markers falling closer to the center). Across cells, phase preference was highly variable in half of the experimental animals (R602, R636). For R887 and R911, theta cell firing converged on a single phase in both gRSC and dRSC. A schematic diagram of results from A and B is shown below each group. A value of 180*^◦^* was assigned to theta troughs and 0/360*^◦^* to peaks, corresponding to opposite hemispheres in the circle.

### 2.0 Retrograde tracer experiment

#### **2.2 Injection coordinates**

The present section provides a summary of all injection coordinates and tracers used. In the tables, s and d indicate whether the center of the tracer deposit in RSC was located in superficial or deep layers (or both). The different AP-levels of the injections are summarized as caudal, rostral or at electrode level in the main text.

**Supplementary Table 3.** List of injection coordinates for thirteen retrograde tracer injections (n=10 animals) targeting RSC at approximately the same AP levels as the recording sites (-5.5 and -5.8 from bregma). Weight refers to weight in grams at the time of surgery.

| ***Injections at electrode level*** | | | |
| --- | --- | --- | --- |
| ***Case No.*** | ***Injection site*** | ***Weight (g)*** | ***Laboratory*** |
| 222#9 | dRSC s + d  (-5.0mm, CTB) | 297 | Aggleton; Cardiff |
| 222#10 | dRSC s + d  (-5.0mm, CTB) | 306 | Aggleton; Cardiff |
| 077941#13 | gRSC d  (-5.5mm, CTB) | 286 | Aggleton; Cardiff |
| 077942#14 | gRSC d  (-5.5mm, CTB) | 227 | Aggleton; Cardiff |
| 878 | dRSC, d+s  (-5.5mm, Ctb-488) | Unknown | Jeffery; London |
| 878 | gRSC, s (d)  (-5.5mm, Ctb-594) | Unknown | Jeffery; London |
| 731 | dRSC, s + d  (-5.5mm, FG) | Unknown | Jeffery; London |
| 731 | gRSC, s + d (  -5.5mm, Ctb-594) | Unknown | Jeffery; London |
| 730 | dRSC, d  (-5.5mm, Ctb-594) | Unknown | Jeffery; London |
| 730 | gRSC s + d  (-5.55mm, FG) | Unknown | Jeffery; London |
| 717 | dRSC/M2 s + d  (-5.55mm, FG) | Unknown | Jeffery; London |
| 729 | dRSC s + d  (-5.5mm, FG) | Unknown | Jeffery; London |
| 682 | gRSC s (d)  (-5.5mm, Ctb-594) | Unknown | Jeffery; London |
| FB, fast blue; FG, fluorogold; CTB/Ctb, cholera toxin subunit b. | | | |

**Supplementary Table 4.** List of injection coordinates for sixteen retrograde tracer injections (n=13 animals) that targeted portions of RSC > 0.5 mm rostral or caudal to the electrophysiology recording sites.

| ***Injections at rostral and caudal locations*** | | | |
| --- | --- | --- | --- |
| ***Case No.*** | ***Injection site*** | ***Weight (g)*** | ***Laboratory*** |
| 187#9 | gRSC d (s)  (Rostral, BDA) | 293 | Aggleton; Cardiff |
| 64#3 | dRSC/M2/V2, s + d  (Rostral + caudal, FB) | 329 | Aggleton; Cardiff |
| 172#27 | dRSC (M2/V2), s + d  (Rostral + caudal, FB) | 295 | Aggleton; Cardiff |
| 172#28 | dRSC (M2/V2), s + d  (Rostral + caudal, FB) | 316 | Aggleton; Cardiff |
| 225#4 | dRSC s + d  (3.30mm/3.80mm, CTB) | 291 | Aggleton; Cardiff |
| 225#12 | dRSC s + d/gRSC d (3.80mm/4.55mm/5.30mm, CTB) | 294 | Aggleton; Cardiff |
| 223#1 | dRSC s + d  (3.50mm/4.50mm, CTB) | 321 | Aggleton; Cardiff |
| 223#4 | dRSC s+d/gRSC d  (4.5mm, CTB) | 329 | Aggleton; Cardiff |
| 227#22 | dRSC s+d  (-3.30, FB) | 463 | Aggleton; Cardiff |
| 227#22 | dRSC d  (-6.80, CTB) | 463 | Aggleton; Cardiff |
| 227#24 | dRSC s + d  (-3.30mm, FB) | 508 | Aggleton; Cardiff |
| 227#24 | gRSC d  (-6.80mm, CTB) | 508 | Aggleton; Cardiff |
| 227#16 | dRSC d  (-3.30mm, CTB) | 606 | Aggleton; Cardiff |
| 227#16 | dRSC s+d/gRSC d  (-6.50mm, FB) | 606 | Aggleton; Cardiff |
| 227#202 | gRSC (dRSC) s + d  (-6.3mm/-6.8mm), CTB) | 519 | Aggleton; Cardiff |
| 227#204 | gRSC s + d  (-6.3mm/-6.8mm), CTB) | 614 | Aggleton; Cardiff |
| FB, fast blue; CTB, cholera toxin subunit b; BDA, biotinylated dextran amine. | | | |

AP levels determined from sections. Where the AP level is extensive, we report whether the injection targeted rostral or caudal sites, without specifying the exact AP of each injection. Weight refers to weight in grams at time of surgery.

#### **2.2 Injection sites for injections made at more rostral and caudal RSC locations**

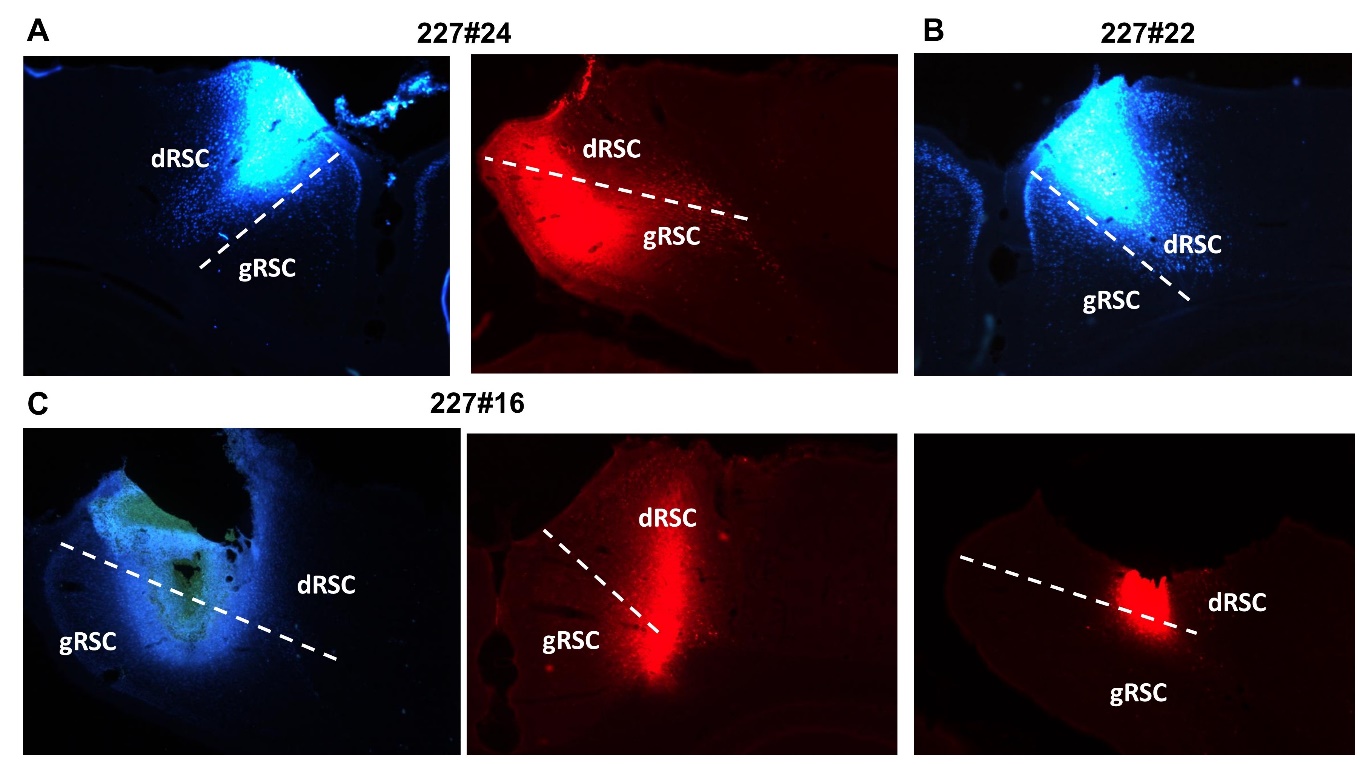

**Supplementary Figure 2. Dual retrograde tracer injections at different AP levels in the RSC**. **(A)** 227#24 Right, caudal CTB injection in gRSC (mainly deep layers); left, rostral fast blue (FB) injection in dRSC (both superficial and deep layers). **(B)** 227#22 Both injections centered in dRSC. Top, rostral FB injection (superficial and deep layers of dRSC); bottom, caudal cholera toxin subunit b (CTB) injection (deep layers of dRSC). **(C)** 227#16 Right, rostral CTB injection mainly confined to dRSC (deep layer); left, caudal FB injection in both dRSC (superficial and deep layers) and gRSC (deep layers).

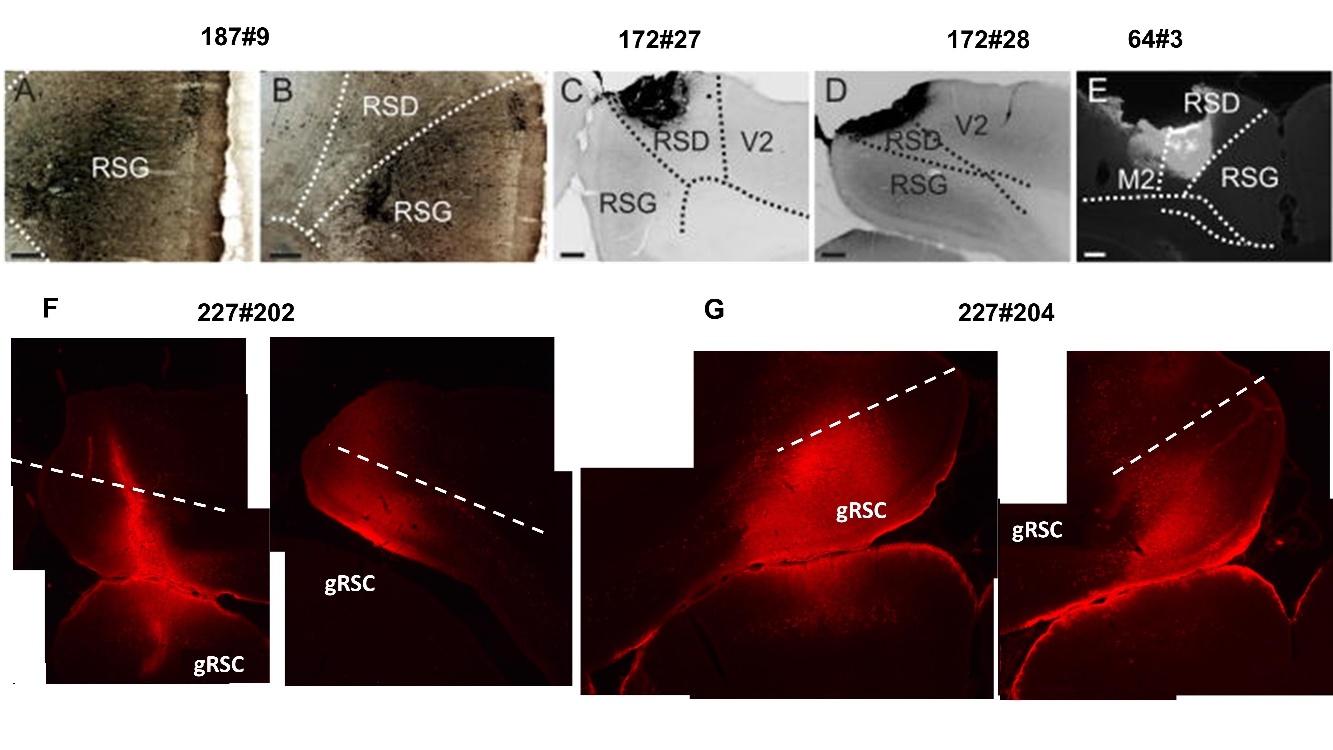

**Supplementary Figure 3. Single retrograde tracer injection in either gRSC or dRSC**. **(A-B)** 187#9 Brightfield images of the biotinylated dextran amine (BDA) injection site centered in gRSC (largely deep), adapted from Mathiasen et al. (2017). **(C-D-E)** Inverted greyscale fluorescence images of FB tracer injection in dRSC, adapted from Mathiasen et al. (2017). All layers seemingly involved**. (F-G)** 227#202, 227#204. Two CTB injection cases in caudal gRSC, all layers (superficial and deep) involved with possible weak dRSC involvement in case #202.
